# The “dark matter” of protein variants carries a distinct DNA signature and predicts damaging variant effects

**DOI:** 10.1101/2021.05.27.445950

**Authors:** Joseph Chi-Fung Ng, Franca Fraternali

## Abstract

Signatures of DNA motifs associated with distinct mutagenic exposures have been defined for somatic variants, but little is known about the consequences different mutational processes pose to the cell, especially how mutagens exert damage on specific proteins and their three-dimensional structures. Here we identify a DNA mutational signature which corresponds to damaging protein variants. We show that this mutational signature is under-sampled in sequencing data from tumour cohorts, constituting the “dark matter” of the mutational landscape which could only be accessed using deep mutational scanning (DMS) data. By training a set of gradient boosting classifiers, we illustrate that DMS data from only a handful (≈ 10) of experiments can accurately predict variant impact, and that DNA mutational signatures embed information about the protein-level impact of variants. We bridge the gap between DNA sequence variations and protein-level consequences, discuss the significance of this signature in informing protein design and molecular principles of protein stability, and clarify the relationship between disease association and the true impact mutations bring to protein function.

## Introduction

High-throughput sequencing methods have enabled the discovery of a large number of genetic variants between individuals, as well as those arisen in disease conditions such as cancers. All somatic cells are vulnerable to genetic variants contributed by various endogenous and exogenous processes. The concept of “mutational signatures” associates each of these mutagenic processes to the DNA alterations it generates, by specifying the sequence contexts proximal to 5’ and 3’ of base substitution events [1, 2]. Algorithms to extract such signatures [1, 2], together with sequencing studies of *in vitro* systems exposed to specific mutagens [3, 4], allow for cataloguing the footprints different mutagenic processes leave on the somatic genome [5]. These studies link various epidemiological risk factors of cancer to their molecular effects at the single-base resolution [2, 6]. Mutational signatures also highlight hitherto under-appreciated endogenous processes which contribute somatic mutations, most notably the APOBEC3 (apolipoprotein B mRNA editing catalytic polypeptide-like 3) enzymes. A large body of work has established the expression pattern of these enzymes [7] and the genomic contexts of APOBEC3 mutagenesis [8], including their preferences in terms of 3D DNA conformational features [9].

While much research has focused on defining the causes of various mutational signatures, the consequences of variants attributable to distinct DNA substitution patterns have not been analysed in a systematic manner. This issue remains challenging to study because existing signature discovery methods do not explicitly assign causal mutagenic process to individual substitutions [2]; a mutational signature is often collation of several DNA motifs; for motifs which are associated with numerous signatures, attribution of causal mutagenic process becomes conceivably more challenging [1, 2]. As a result, existing methods which annotate consequences of genetic variants and predict cancer driver genes have yet to systematically identify the responsible “culprit” of specific damaging mutations. This, however, could have great significance and benefit to advance our understanding of cancer biology, in rationalising how lifestyle and clinicopathological features relate to damages in our genomes, linking the process of mutagenesis to their effects in gene products, and monitoring the acquisition of mutations attributable to drug regimens that could present as side-effects in cancer patients who undertake these treatments. From a basic science perspective, a signature of protein damage also offers a route to identify attractive targets for protein design (to remove “fragile points” that can be easily destabilised from the protein), and more broadly informs the molecular principle of protein stability.

There are important considerations in the approach to comprehensively survey the impact of variants. Notably, biological samples are constantly under evolutionary selection pressure which removes genetic variants that are deemed harmful to cell viability [10]. In somatic evolution, those variants which enhance viability are retained and positively selected; such signals have been exploited to identify cancer driver genes [11]. Conceivably, one of the correlates for this selection pattern is the localisation of mutations with respect to a protein’s three-dimensional structure, which would imply varying consequences to the expressed polypeptide [12, 13]. We and others have discussed patterns of variant distribution across regions along the protein sequence defined using protein three-dimensional structural information, mapping datasets of variants from health and disease (e.g. [12, 14, 15, 16]), which reveal these consistent patterns: (i) depletion of core variants except for cases where they may be pathogenic (e.g. loss-of-function cancer drivers) and (ii) enrichment of disease-causing variants in the protein interaction interfaces. Sequencing experiments performed on biological samples may therefore be biased by the evolutionary selection acted upon the consequences genetic variants bring to cells: mutational processes which generate the most damaging variants - should such processes be active in the cell in the first place - would be difficult to sample. Such variants could be understood as part of what we could term as the “dark matter” of the somatic mutational landscape, arising from the under-sampling of such variants in observational studies, and therefore remain difficult to study. One way to address this issue is to perform “Deep Mutational Scanning” (DMS) [17, 18] where every position in the protein is mutated to any other 19 amino acids either *in vitro* or *in silico*, followed by assays to measure the stability and/or activity of the mutants. Experimental DMS have been applied on several cancer-related proteins and domains [19, 20, 21, 22]; computationally, variant impact predictors which incorporate sequence conservation (large protein/domain multiple sequence alignments), structural/physicochemical features (protein secondary/tertiary structures) and consequential representations of protein motions (imposing network models onto three-dimensional structures to account for molecular vibrations) have been developed, applied and evaluated in a DMS context [23, 24, 25, 26]. These computational and experimental data allows us to probe and access features of variants which reside in this mutational “dark matter”. To link variant impact to DNA mutational signatures one would require mapping of proteins to their corresponding coding sequences; to this date, such analyses which identify damaging mutational signatures have yet to be performed.

Here we extract a DNA mutational signature, dominated by the motif `5’-G/C[T>G]A/G-3’`, which characterises variants that alters physicochemical properties at protein cores (hereafter ‘damaging protein variants’). We show that these short DNA motifs are under-sampled in mutation data obtained from tumour samples, and could only be accessed using a DMS approach. We train predictors using experimental and computational DMS data, and show that such methods, learning even from only a handful (≈ 10) proteins, are capable of accurately predicting damaging variants for proteins unseen in training, comparable to other variant impact prediction tools. Using these models, we illustrate that DNA signatures, in combination with contextual information (sequence conservation, protein structural environment etc.) of the mutated position, can accurately annotate variant impact. These analyses probe the space of the mutational “dark matter”, and enrich pathogenicity prediction offered by conventional variant impact classifiers which are typically trained without access to these under-sampled but protein-damaging variants.

## Results

### Short DNA motifs are informative in distinguishing damaging protein variants

We and others have reported the distribution of variants across different regions of the protein three-dimensional structures, which points to a consistent depletion of variants in the protein core (i.e. buried from solvent) except for disease-causing variants [12, 14, 15, 16]. Since these studies typically rely on variants observed in biological samples, we reason that this reflects an under-sampling of mutations which are less tolerable to the cell, e.g. by destabilising the implicated proteins (Figure 1A). Different physicochemical environments in protein three-dimensional structures constrain the allowed types of amino acids; this implies a biased distribution of DNA motifs in the coding sequences which, if mutated, would target distinct regions of the protein structure. We therefore hypothesise that there are specific DNA sequence patterns which implicate a damaging protein-level impact. Discovery such DNA motifs requires comprehensive measurements which consider every amino acid positions and survey substitutions to every other 19 amino acids. Such deep mutational scanning (DMS) data [17, 19, 27] represent a suitable data source for unbiased discovery of mutational signatures associated with damaging protein variants. We first obtained experimental DMS data from a public database (MaveDB [27]) on two proteins, PTEN [19, 20] and MSH2 [22]; these are two human proteins harbouring known, missense, cancer driver variants with DMS data available in the database. Noticing the limited availability of experimental DMS data, we further obtained analogous computational DMS data by applying variant impact predictors (PolyPhen-2 [28], EVmutation [29] and rhapsody [24]) over full-length coding sequences of a total of 6,918 human proteins (see Methods). Comparing to the experimental DMS profiles above, this computational dataset greatly expands the extent of the proteome surveyed; moreover, the predictors we used consider different features [24, 28, 29, 30] (sequence conservation, static protein secondary/tertiary structures, and dynamics models defined on structures), taking into account different aspects of the molecular profile in variant impact annotation. We developed a R package (CDSMutSig) with utilities to map any amino acid substitution to the corresponding coding sequence change and DNA mutational contexts (see Methods). We took into consideration the fact that a missense amino acid change could be caused by nucleotide substitution at any position of the codon, depending on the exact missense change in question. Therefore, the trinucleotide mutational contexts could be defined “in-frame” within the same codon, or span tandem codons. Utilities in the CDSMutSig package consider any possible nucleotide (and therefore amino acid) substitution, and allow for filtering to remove amino acid changes which requires tandem nucleotide changes. We evaluate the association of each possible three-base DNA mutational context with mutational impact by fitting linear models of the obtained DMS profiles using trinucleotide mutational signatures as the explanatory variable. These coefficients over all individual DMS models are depicted in Figure 1B. Comparing the regression coefficients, we find that four sequence contexts, namely `G[T>G]G, G[T>G]A, C[T>G]G and C[T>G]A` (these motifs are written in the direction from 5’ to 3’) are associated with damaging variants. These motifs define a DNA mutational signature of damaging protein variants, carrying characteristic nucleotide contexts which can be used to scan the protein-coding genome to prioritise targets for protein design, potentially rescuing these proteins from being endangered by “fragile points” which perturb protein function (see Discussion).

**Figure 1:**
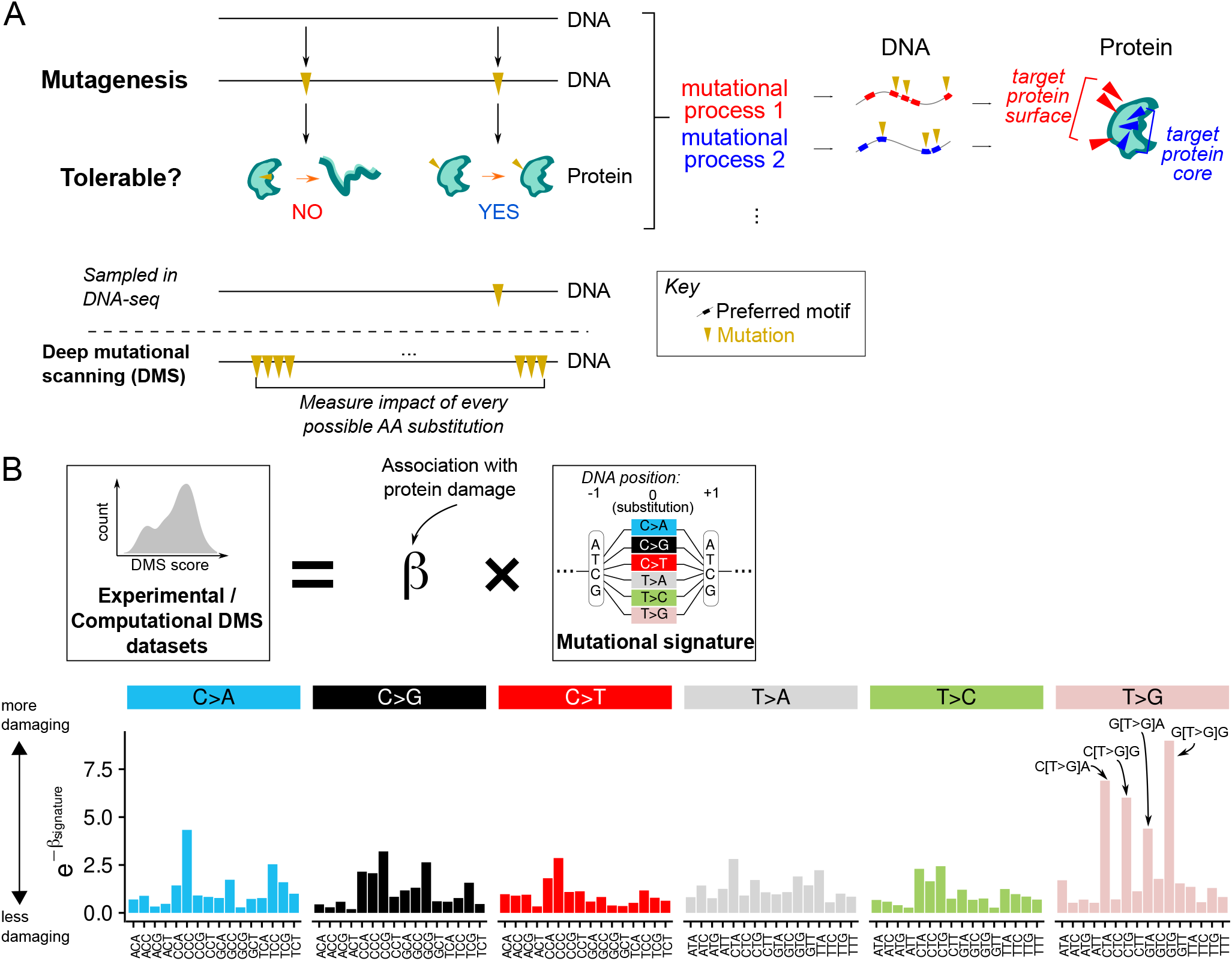
DNA motifs associated with damaging protein variants. (A) Schematic of the overaching hypothesis. Tolerability of mutations at the protein level determines the ability to sample such mutations in conventional DNA sequencing experiments. Different mutational processes prefer distinct motifs that may display preference to localise to specific protein regions (right, here illustrated with protein surface and core as examples), due to amino acid preference in different protein three-dimensional structural environments. Deep mutational scanning (DMS) overcomes the under-sampling of intolerable variants by exhaustively measuring impact of every possible amino acid substitutions on the protein. (B) To identify mutational signatures associated with damaging variants, we fit linear models on experimental and computational DMS datasets, using mutational signature (expressed as trinucleotide contexts surrounding the nucleotide substitution) as covariate. The coefficients *β* indicates the association of a given trinucleotide context to damaging DMS scores. This was performed on *n* = 6 experimental/computational DMS screens, and the mean of the coefficients for each trinucleotide motif (*β*_signature_) was depicted as a bar plot. This mean is transformed as a negative exponent (expressed as 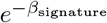 in the axis label for simplicity) such that contexts which are more strongly associated with damaging protein variants are signified by higher bars.

Next we investigate the biophysical basis behind this damaging mutational signature. Figure 2A shows the amino acid substitutions attributable to each of the damaging mutational contexts, had such mutations occurred and been sampled in sequencing experiments. We observe that these mutational contexts bring about three main alterations to the physicochemical properties of the mutated amino acid positions (Figure 2A): (i) change in charge (mutations to arginine/aspartate); (ii) change in polarity (e.g. tyrosine → serine); change in structural flexibility (mutations to glycine/proline). We further find that all such contexts tend to localise to protein cores relative to surface and interface (Figure 2B). Together, these results provide biophysical explanations, at the protein level, behind damaging mutational impact associated with specific DNA sequence contexts. We demonstrate that the use of DMS data can offer opportunities to discover damaging variants which are difficult to sample in observational studies.

**Figure 2:**
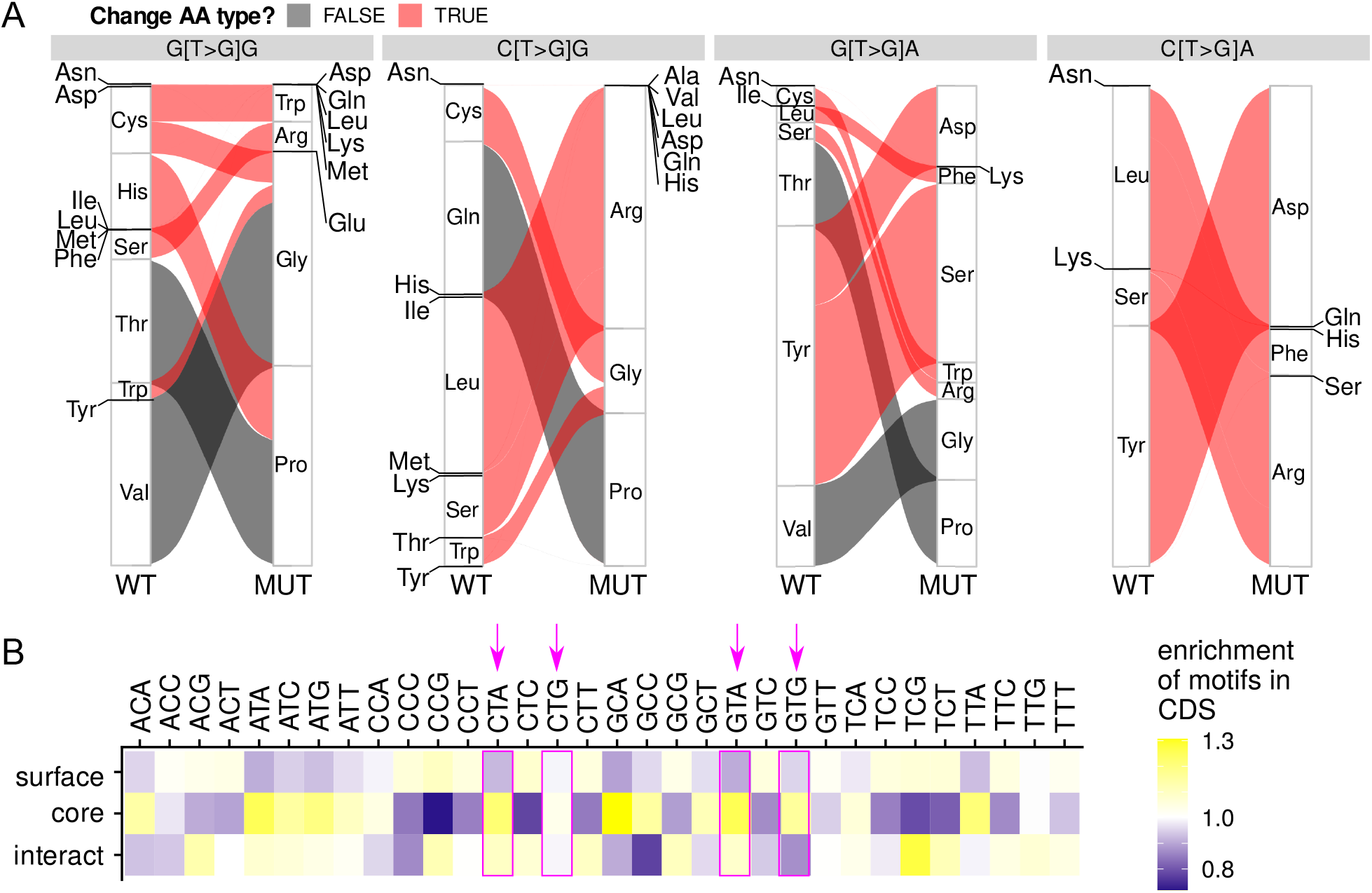
Physicochemical characterisation of the damaging mutational signature. (A) Amino acid (AA) substitutions attributed to each damaging mutational context highlighted in Figure 1B. The width of ribbons correspond to the amount of AA substitutions assayed in the computational DMS screen performed using rhapsody [24]. Ribbons are coloured to indicate whether the AA change constitute changes in physicochemical properties. (B) The enrichment of mutational motifs in protein surface, core or interface calculated over all human coding sequences (CDS) with structural coverage. The motifs highlighted in the damaging mutational signature are highlighted in magenta.

### Observed somatic mutational profiles under-sample damaging protein variants

We further characterise this under-sampling of damaging variants in sequencing studies of tumour cohorts. Overlaying ClinVar (which represents variants with clinical evidence of pathogenicity) and TCGA (The Cancer Genome Atlas, i.e. variants discovered in whole-exome sequencing of tumour samples) variants onto the experimental DMS datasets considered in the previous analysis, we see that TCGA variants have less severe mutational impact than known pathogenic variants (Figure 3A), suggesting TCGA data are dominated by signals from passenger variants with little impact. To compare the effect of mutations which dominate the observed somatic mutation landscape against the damaging mutational signature described above, we further select DNA sequence motifs which represent six common mutagenic processes as case studies (Figure 3B). These cover well-studied endogenous and exogenous mutagenic processes (Aging, APOBEC3, ultraviolet light [UV], and hypermutation caused by *POLE* mutations) [1] as well as common therapeutic strategies applied in cancer treatment (5’-fluorouracil [5-FU] and Platinum-based therapies) [5]. Each of these mutational signatures can be attributed to distinct recurrent variants observed in different tumour types (see Supplementary Note 1). In contrast with the damaging mutational signatures, which shows a consistent association with a damaging score across all examined DMS datasets, these common signatures display much weaker association with damaging protein impact (Figure 3C). We also observe that in large tumour cohorts such as TCGA [31], whilst the common cancer signatures are well-represented, the damaging sequence contexts are much more rarely mutated (Figure 3C). This suggests that the damaging signature, unlike the common signatures which are driven by active carcinogenic processes, is either inactive in a cancer context, or are under-sampled in the data due to selection effects. We then compare the general trends of TCGA mutations that occur in contexts attributable to each of the signatures in terms of their localisation to protein surface, core and interface. For all six mutational signatures, an enrichment for mutations on protein surface and depletion in the protein core and interacting interface can be observed (Figure 3D). However, these regions do not show large differences in terms of the amount of mutable DNA motifs in the native coding sequences (Figure 3E). These results support the dominance of tolerable mutations in observed mutational profiles (which could be further confirmed by more detailed analysis of the selection patterns in different protein structural regions, see Supplementary Note 2), and suggest that common, active mutagenic processes avoid DNA motifs which could give rise to damaging amino acid substitutions.

**Figure 3:**
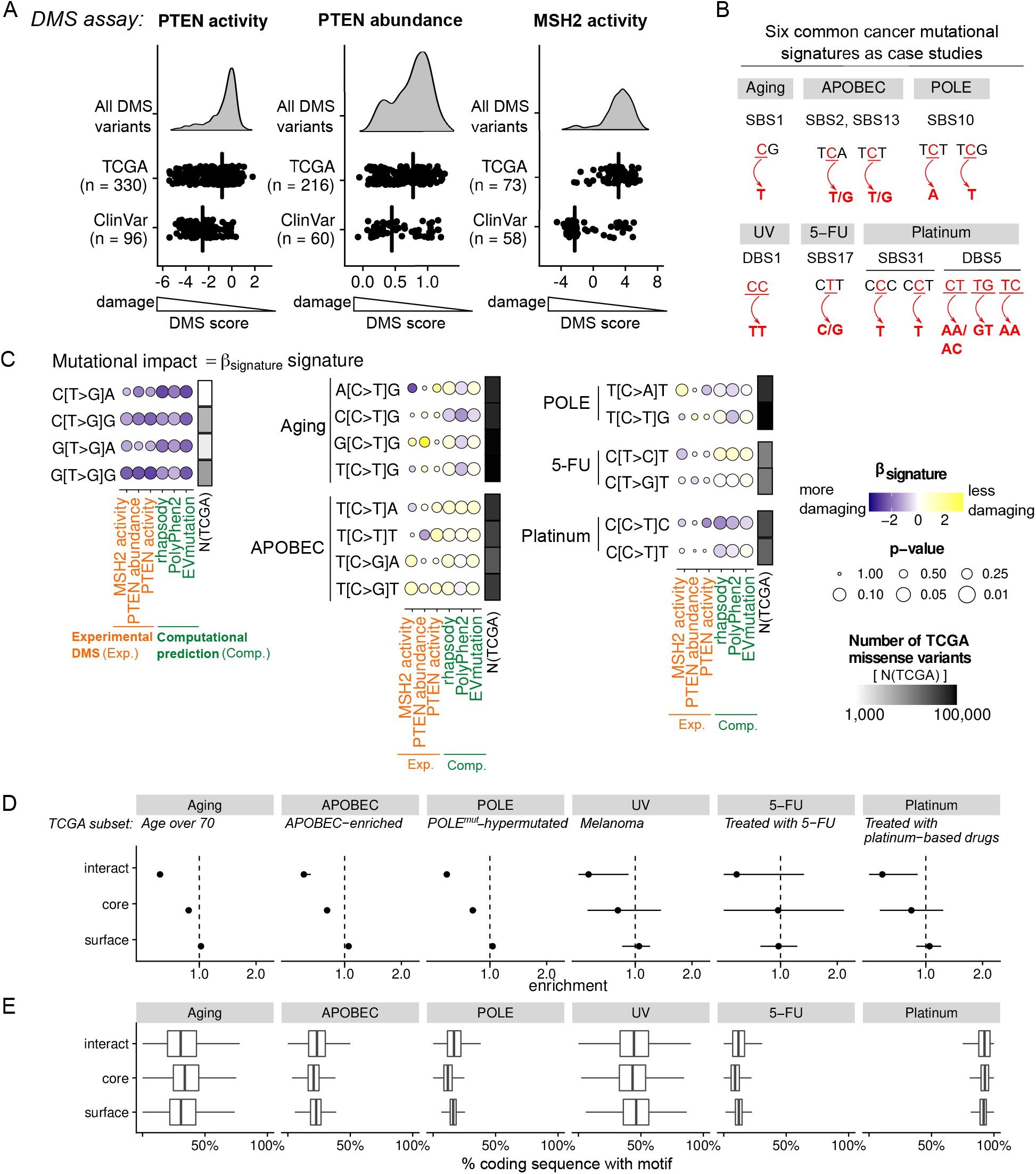
Observed somatic mutational profiles under-sample damaging protein variants. (A) Experimental DMS profiles of two cancer-related proteins, PTEN and MSH2, obtained from MaveDB [27]. The impact scores of ClinVar and TCGA variants are depicted as dotplots. Vertical bars indicate medians. (B) Six mutational signatures common in somatic mutations are considered as case studies. The preferred DNA motifs in each case are depicted. SBS, single-base substitution; DBS, doublet-base substitution. The nomenclature of mutational signatures according to the COSMIC catalogue (v3.1, https://cancer.sanger.ac.uk/cosmic/signatures/) are noted. (C) The association of trinucleotide contexts (vertical axis) to experimental or computational DMS scores (horizontal axis) is represented by the regression coefficients *β*_signature_ (summarised in Figure 1B). Here the contexts for the damaging variants (left column) are compared against those represented in the common cancer signatures (right). The number of TCGA variants occuring in each context is depicted in greyscale (rectangles next to the *β*_signature_ dot plot). See Supplementary Figure S3 for the complete plot, and Supplementary Table S4 for data underlying this analysis. (D) Variant enrichment in protein structural regions for the six mutational signatures in different defined subsets of the TCGA data (indicated in italics). Error bars depict 95% confidence interval estimated by bootstrapping. (E) Distributions of the abundance of motifs (in terms of the percentage of human coding sequences with such motifs) in native, wild-type coding sequences associated with each signature in protein surface, core and interacting interface.

### DMS data can be utilised for probing the mutational “dark matter” and variant impact prediction

With the mutational signature for damaging protein variants being under-sampled in large-scale mutation databases and only accessible by analysing DMS data, we ask whether DMS can be used further to define and probe the scope of such under-sampled protein variants (i.e. the mutational “dark matter”). We reason that protein variants can be segregated by two levels of impact: (i) pathogenicity, i.e. their association to disease onset and/or development, and; (ii) damage to the stability and/or activity of proteins. The “dark matter” would correspond to variants which are highly damaging to protein stability/activity, but without evidence of pathogenic associations (Figure 4A). Using these two variables we can also define other types of variants: polymorphisms (neither pathogenic or protein-damaging), gain-of-function (GOF; pathogenic but not protein-damaging). The final category (pathogenic and protein-damaging) appears problematic at first sight; this corresponds to variants which causes damage to proteins, but can be offset by the resulting selective advantage towards cell viability; we term these as loss-of-function (LOF) variants (Figure 4A). To test this idea we select two *in silico* scoring systems trained to predict these two variables: first, the boostDM [26] algorithm distinguishes cancer drivers from non-drivers, representing the pathogenicity axis of this landscape. Second, the EVmutation algorithm [29] takes into account epistatic interactions between protein residues and fits a statistical potential to quantify mutational impact based on a multiple sequence alignment of homologous domains; we reason that this could be a good proxy for protein-level damage (Figure 4A). We consider all protein variants screened using these two *in silico* scoring systems and plot the distribution of variants in this space. Figures 4B and C show the distribution of PTEN and TP53 protein variants by the boostDM and EVmutation scores, separated by their localisation in protein structural regions. Whilst the shapes of these distributions appear to vary from protein to protein, we see that surface variants tend to be mapped to polymorphisms, and core variants to the LOF and “dark matter” regimes. An enumeration of variants mapped to each of the quadrants in the schematic in Figure 4A for different mutation databases confirm the under-sampling of “dark matter” variants in all databases in comparison to DMS; the mutational “dark matter” itself appears to be enriched in core variants (Figure 4D). Taken together, our analysis defines and links properties of damaging protein variants: they carry a distinct DNA motif, introduce specific amino acid changes, target buried regions in protein structures, and constitute the bulk of the mutational “dark matter”.

**Figure 4:**
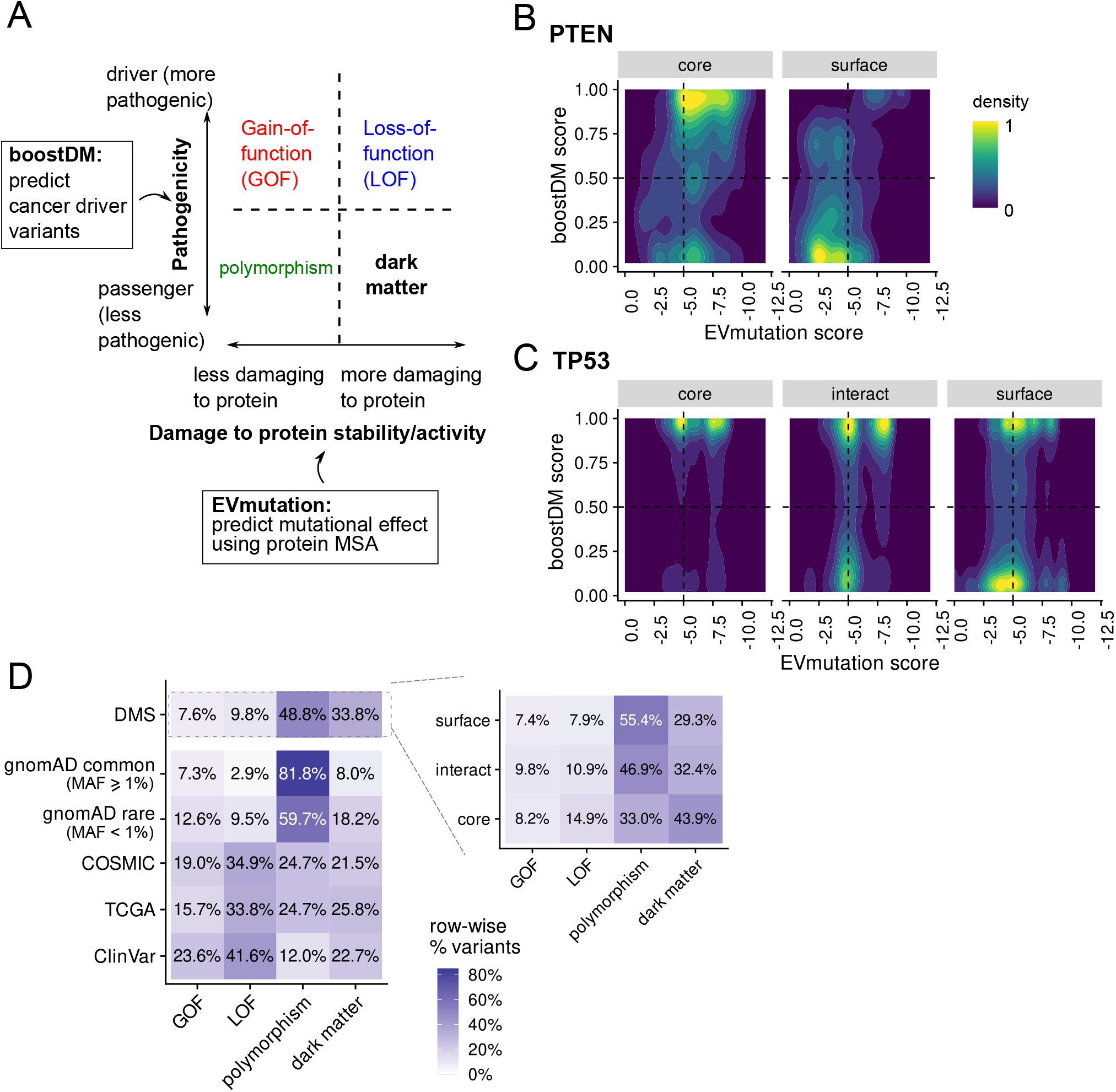
Defining mutational “dark matter” and the variant impact landscape using *in silico* DMS predictors. (A) Schematic of the variant impact landscape defined by variant pathogenicity and its damage to protein stability/activity. boostDM [26] and EVmutation [29] scores are used as proxies for these two variables. (B,C) Distribution of all possible amino acid variants in PTEN (panel B) and TP53 (C) with respect to their boostDM and EVmutation scores. Variants are separated by their protein structural localisation. (D) Statistics of variants in the gain-of-function (GOF), loss-of-function (LOF), polymorphism and the MDM (“dark matter”) quadrants of the landscape depicted in panels A-C. The proportions of variants in these categories are quantified for the ClinVar, TCGA, COSMIC and gnomAD (further separated into common [minor allele frequency, or MAF ≥ 1%] and rare subsets) databases, as well as all possible amino acid variants (i.e. DMS) across all *n* = 49 proteins with available boostDM and EVmutation scores. Inset shows the breakdown of variants across protein structural regions for the DMS data.

DMS offers access to the mutational “dark matter” unrivaled by large mutational databases; with its exhaustive nature of surveying every possible protein variants we hypothesise that DMS represents a rich resource of training data for variant impact prediction, upon rational selection of mutation features (e.g. solvent accessibility as discussed above) which robustly segregate variant impact. Using experimental DMS data from MaveDB [27], we construct a series of gradient boosting classifiers to evaluate the potential of variant impact prediction using DMS data. Gradient boosting classifier is chosen both because of its capability for feature interpretation and its performance in previously reported variant impact prediction tasks (e.g. boostDM [26]). We frame this as a binary classification task by converting the variant impact scores obtained in individual DMS experiments into binary (damaging/not damaging) labels, and annotating variants with four features: (1) wild-type and mutant amino acids; (2) trinucleotide motif surrounding the DNA substitution (hereafter referred to as DNA mutational signature); (3) conservation of each position of the trinucleotide motif (using PhyloP [34]); (4) solvent accessibility (the quotient solvent accesisble surface area, or Q(SASA), calculated using POPS [35]; see Methods for detailed description of these metrics). In contrast to previous machine learning approaches utilising DMS data (e.g. [36, 37, 38]), here we introduce DNA mutational signatures as a feature to be evaluated for its predictive utility, and explicitly test the contribution of different feature combinations in classifying variant impact (Figure 5A). We note that only a small number (*n* = 29 proteins from 83 experiments) of experimental DMS datasets are publicly available (as of August 2021), with even fewer proteins (9 proteins from 10 experiments) displaying a bimodal distribution of DMS scores amenable for a binary clasification task. We therefore build analogous variant impact predictors using the *in silico* EVmutation [29] scores as the training data including DMS predictions covering 6,671 proteins (Figure 5A). Since EVmutation deploys a statistical potential modelled on protein multiple sequence alignments, we choose it over the other computational scoring systems (boostDM, rhapsody etc.) to avoid accumulating errors over results offered by different machine learning architectures. Comparing with databases which collect *in vitro* thermal stability data for protein variants (e.g. ProThermDB [32] and ThermoMutDB [33]), the experimental DMS dataset which we curate here (hereafter DMSexp) already provides more variants to train classification algorithms (Figure 5B), even though the number of proteins covered is highly limited. Overall, these variant impact classifiers perform well, with a receiver operating characteristics area-under-curve (ROC-AUC) of up to 0.868 (Figure 5C) when tested on variants withheld from training. We compare our predictions against Envision [36], a machine learning method also trained on experimental DMS data, and find that our model reproduces the experimental scores to a comparable, and in some case superior, accuracy than Envision (Figure 6A). We note, however, that Envision frames the problem as a regression task (i.e. deriving a score to measure variant damage) whereas our DMSexp prediction treats this as a classification problem, optimising not for correlations with experimental DMS scores but for the accuracy of a binary classification. This potentially explains the modest associations with DMS scores; the availability of more experimental DMS datasets in the future would allow more extensive benchmarking of these different predictors.

**Figure 5:**
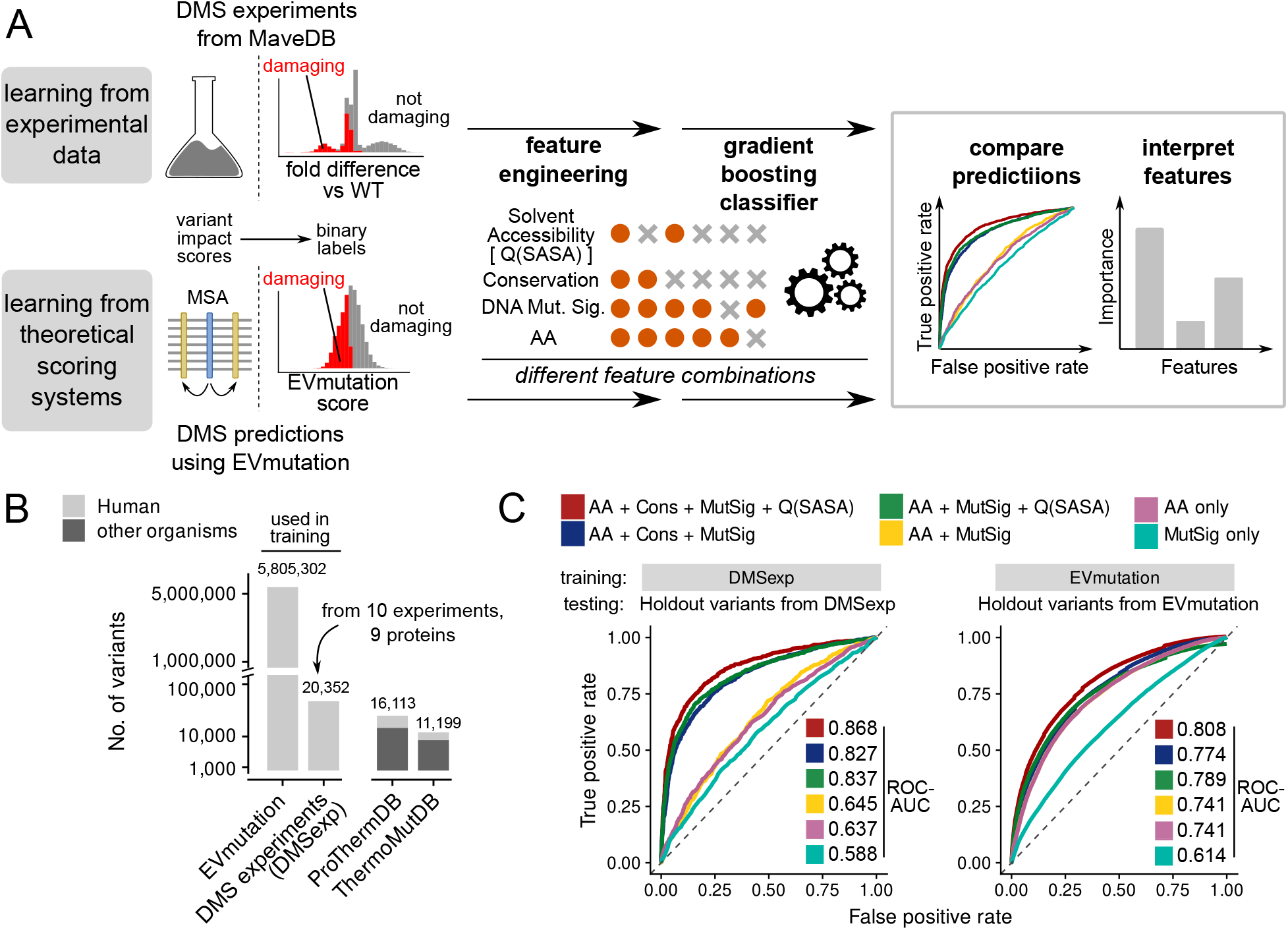
Using DMS data to predict variant impact. (A) Schematic of the workflow to construct gradient boosting classifiers using experimental (from MaveDB [27]) and theoretical (EVmutation [29]) DMS data to predict the damaging impact of variants. (B) The number of variants covered in EVmutation and DMS experiments (“DMSexp”) used in training the classifiers. The numbers of variants documented in ProThermDB [32] and ThermoMutDB [33] (as of December 2021) are included for comparison. Supplementary Table S1 contains the underlying numbers. (C) Receiver operating characteristic (ROC) curves for classifiers trained on DMSexp and EVmutation data using different feature combinations, tested on variants from the respective dataset but withheld from training. The area-under-curve of these ROC curves (ROC-AUC) are indicated.

**Figure 6:**
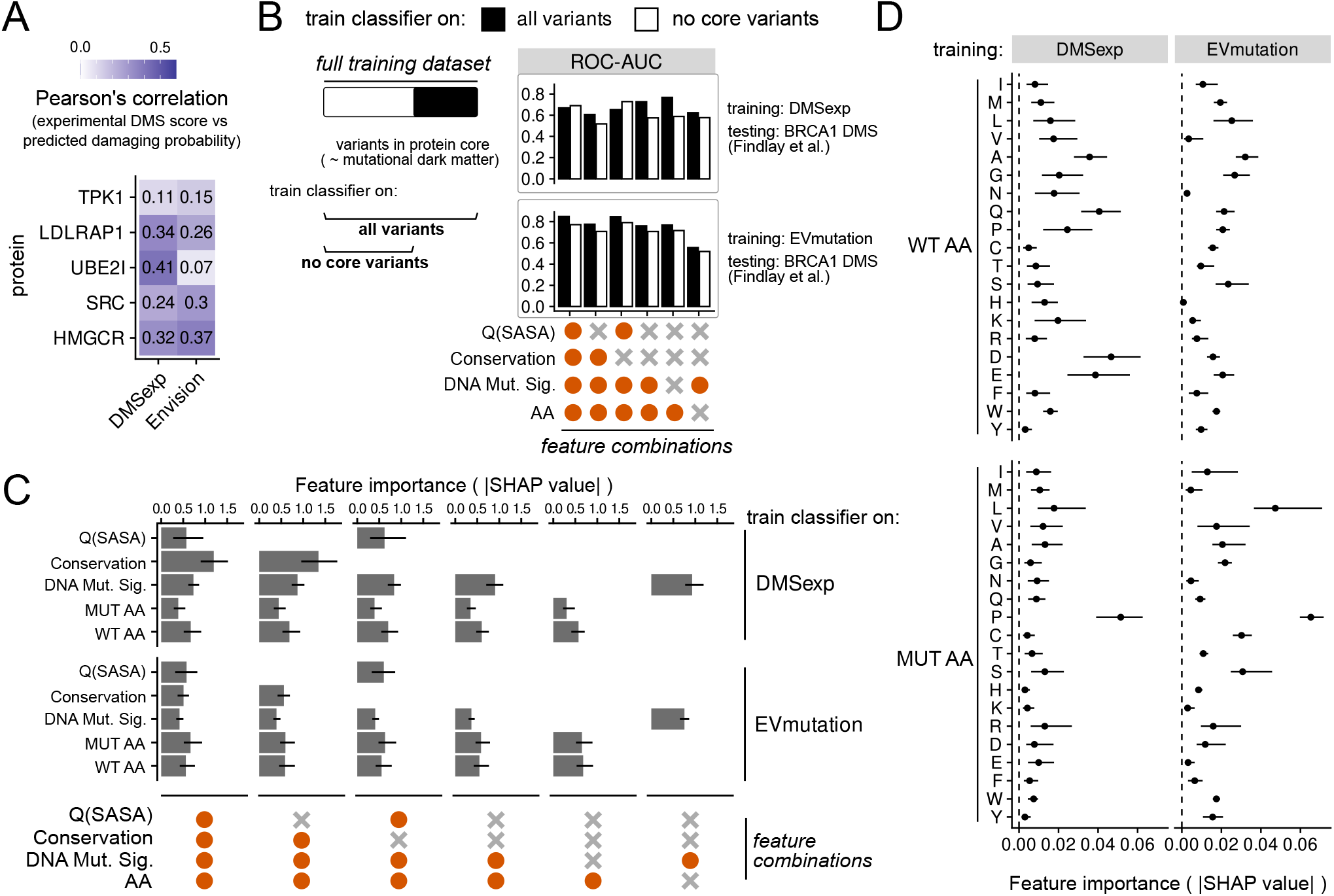
Interpreting variant impact predictors trained on DMS data. (A) Predictions offered by the DMSexp model are compared against experimental DMS scores obtained for *n* = 5 proteins, displayed as the Pearson’s correlation. This is compared against analogous prediction offered by Envision [36]. (B) Comparison of performance of predictors trained on the entire dataset versus those where variants in the protein core (a proxy for mutational “dark matter’, see Figure 4) are removed. The predictors are tested on an experimental DMS dataset on the BRCA1 protein [21]. The ROC-AUC metrics are depicted. Detailed statistics are given in Supplementary Table S3 and Supplementary Figures S4-S6. (C) Feature importance (quantified using the Shapley additive explanations [SHAP] method [39]) quantified on the classifiers trained on DMSexp (top) and EVmutation (bottom) datasets. Grey bars depict the median absolute SHAP value (the higher this value, the higher its contributions towards the predictions) and error bars depict the inter-quartile range over all predictions on holdout variants from the respective datasets. (D) Feature importance broken down by individual wild-type (WT) and mutant (MUT) amino acids. Dots depict the median and error bars depict the inter-quartile range of all predictions on holdout variants from DMSexp (left) and EVmutation (right) datasets.

We next assess whether these predictors can infer variant impact for proteins unseen from training, and specifically ask whether the unique access to the mutational “dark matter” offered by DMS data improves variant impact predictions. Considering that the “dark matter” is enriched with core variants (Figure 4), we remove core variants from the training data and re-train variant impact predictors. We test these predictors on a DMS dataset performed on the BRCA1 protein [21] which is unseen during training. On this task, the predictors display a similar level of performance (ROC-AUC ≈ up to 0.8), with the “full” versions surpassing the predictors trained without access to the core variants as a proxy to the mutational dark matter in all tested feature combinations (Figure 6B); the only exceptions are in cases where the predictor relies directly on solvent accessibility information to classify variants. These data suggest that DMS, and specifically the mutational dark matter, are useful as a resource for variant impact prediction. In particular, considering the restricted scope of proteins included in the experimental DMS data, our results suggest that these limited data already embed sufficient information for robust and accurate annotation of variant impact.

We further analyse these predictors to assess whether DNA mutational signatures contribute to the predictions. We note that while mutational signature alone constitutes a predictor which is only slightly better than random (Figure 5C), it bears importance comparable to other features when used together for training (Figure 6C), especially in predictors trained on the DMSexp dataset, suggesting that DNA mutational signatures embed information that is useful for variant impact prediction in conjunction with other mutational features. We also observe that mutations involving specific amino acids, especially substitutions to proline, are particularly revealing of variant impact (Figure 6D). Taken together, these analyses reveal how different mutational features could contribute to accurately annotate variant impact, and suggest that a computational approach which benefit from DMS analyses could learn the general features governing variant impact and perform inferences for proteins and variants not yet covered with DMS experimental efforts.

## Discussion

In this study we identify a DNA mutational signature associated with damaging protein variants using DMS data, and demonstrate the use of DMS in probing mutational “dark matter” and offering predictive utility for variant impact. To our knowledge this is the first work defining a mutational signature which signifies the consequences of mutations at the protein level, as opposed to those characterised by a given mutagenic cause. The exact genomic locations mutated upon exposure of a given mutagenic process are known to vary across different biological samples and experimental conditions [40], therefore it is impossible to attribute, with certainty, mutagenic processes with individual genetic variants. Our case studies (Figure 3) of signatures which are each over-represented with variants occurring within specific DNA motifs illustrate the likely impact these signatures could cause to individual proteins. For many of these common mutational signatures, we are able to observe, from sequenced tumour samples, recurrent destabilising variants such as those residing in the cancer drivers PTEN [41] and PIK3CA [42], the causes of which have previously been linked to the respective mutagenic exposures. Our mutational signature of damaging protein variants, on the other hand, describes DNA motifs which, if mutated, would likely render damage to protein stability and/or function. Whilst the fact that this signature is only accessible in DMS data but rarely sampled in observed mutational profiles (Figure 3) precludes analyses of the activity of such mutagenic process in a cellular context, this signature for damaging protein variants offers rational guidance to select targets in protein design and therapeutics. By scanning the protein-coding genome for such nucleotide contexts, proteins with such DNA motifs at the coding DNA sequence of important functional sites represent targets which can potentially be easily destabilised to lose its function. These are ideal targets for design with the aim to “rescue” such proteins from the vulnerability of damage, by removing these “fragile points” at their coding sequences. Knowledge about the relationship between nucleotide contexts and protein damage could also inform the design of protein therapeutics, including antibodies, to improve their stability and other biophysical properties.

The most important finding is that DNA motifs as short as trinucleotides embed information which are indicative of variant impact at the amino acid level. The DNA mutational signature we extracted (Figure 1B) features highly specific trinucleotide motifs which can be directly linked to dramatic alterations of physicochemical properties at the protein core (Figure 2). Residue solvent accessibility, as well as other protein structural features, have been heavily studied in terms of their utility in variant impact annotation [12, 14, 15, 16], and in protein evolution to explain differences in evolutionary rates across different protein regions [43]. Here we show additionally that in conjunction with other protein-level features, it holds predictive power for variants without known variant impact annotation when we train classifiers based on these features using DMS data (Figures 5,6). We note that DMS datasets have previously been utilised to construct variant impact predictors with reasonable success (e.g. [25, 36, 37, 38]); here our work serves the additional purpose of validating the relevance of DNA signatures in annotating variant impact. It is also remarkable that the damaging mutational signature is visible only in the analysis of DMS data, but under-represented in mutational profiles obtained from tumour samples. The finding that very few mutations that occur in such contexts can be observed in TCGA (Figure 1B) could either suggest that (a) an active mutagenic process which is capable to generate mutations occurring in these DNA motifs is absent in human cancers, or; (b) these mutations bring excessive damage to protein stability/activity, without sufficient compensatory proliferative advantage to tumour evolution. It is difficult to ascertain which explanation is more plausible. The advancement in the sensitivity to detect ultra-rare variants in the genome [44] might help in determining whether this signature is biologically active at all. The damaging nucleotide contexts could potentially represent a therapeutic opportunity if mutators specific to these DNA motifs can be designed and directed specifically towards tumour cells; this can orchestrate an increase in tumour mutational burden to accumulate a large amount of damaging genetic variants [45] and constitutes a “killing strategy” for tumours which otherwise do not harbour any molecular alterations targetable using existing therapies.

Our comparison of observed somatic mutations against DMS data illustrate the bias towards tolerable variants in observational studies, and conversely the utility of DMS data in defining the scope of the mutational “dark matter”. With boostDM and EVmutation, we demonstrate that by applying predictors developed using orthogonal information in a DMS fashion, one can attempt to segregate variant pathogenicity from their protein-level damage (Figure 4). We also show that DMS data can provide a comprehensive training dataset to develop predictors sensitive enough to distinguish between intolerable versus tolerable variants (Figure 5,6), where the contributions of different mutational features on predictive accuracy can be compared and evaluated. In contrast, observed mutational profiles are naturally biased towards tolerable mutations (and, in the case of somatic mutations, favourable mutations which bring about proliferative advantage to the cell), while truly damaging mutations (which are more likely removed through tumour evolution) are under-represented. This has important implications in the quest of comprehensive characterisations of variant impact, since damaging variants constitute the mutational “dark matter” which is likely inaccessible using solely sequencing experiments performed on biological samples (Figure 7). The only exception where this under-sampling problem can be avoided is when these variants bring greater proliferative advantage to the cell, that overrides any costs imparted on the stability of the affected protein (Figure 7), e.g. loss-of-function mutations that target the protein cores of tumour suppressor genes [12], which are likely to be positively selected during tumour evolution (Supplementary Note 2). Experimentally, to override the under-sampling problem of damaging variants, one potential solution is to use dedicated methods such as ultra-deep sequencing [46] or emerging single-nucleus sequencing approaches [44] to generate mutational profiles from biological samples. Another potential solution is to generate and analyse DMS data to explicitly sample this space in the variant landscape. DMS has been a rich resource for predicting the three-dimensional folds of protein without resolved structures [47] and designing compensatory mutations which negate effects of a pathogenic variant [48], both of which are of immense interests for molecular biologists interested in the effects of mutations on protein stability and activity.

**Figure 7:**
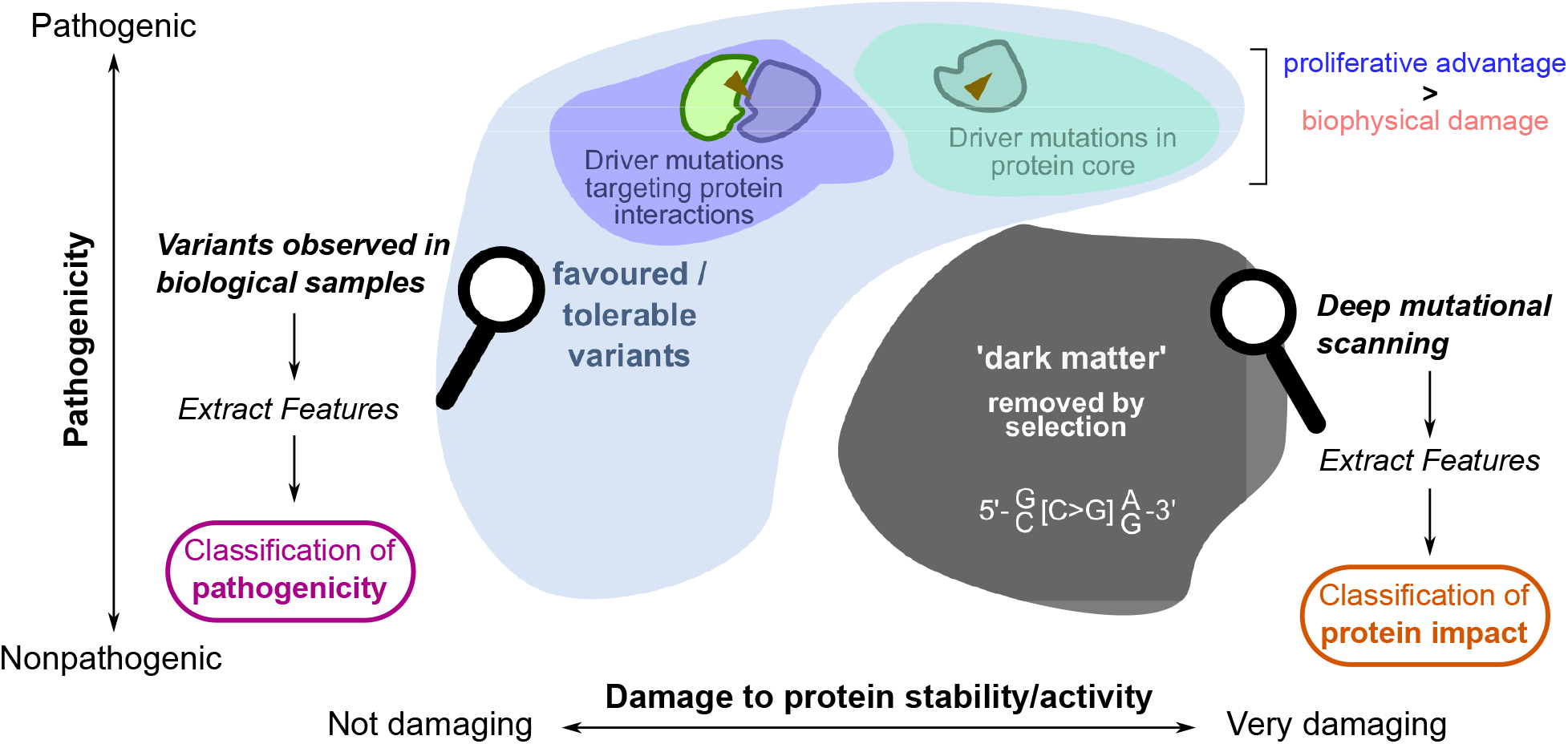
Overview of this analysis. Mutations can be contextualised in terms of their pathogenicity and their impact to the stability/activity of the affected protein. A regime of damaging variant impact constitutes what we term the “dark matter” of somatic mutational landscape which can be characterised by a distinct DNA mutational signature. This regime is under-sampled in sequencing experiments of biological samples, due to the fact that they are likely to be removed by selection. DMS data allow for probing of this space to extract features that explain damage to protein stability and/or activity. Outside of this “dark matter” regime variants are either favoured or allowed, and map to different protein structural regions. Those variants which bring about greater proliferative advantage to offset any biophysical damage will be retained and/or positively selected, as in the case of driver variants targeting protein core and interactions. Such variants can readily be sampled by sequencing, allowing for the discovery of features which explain variant pathogenicity.

The bias towards tolerable variants in observational studies also has important implications in understanding mutational impact and developing variant impact annotation tools. A large number of sequence- and structure-based predictors have been shown to be valuable in offering insights of variant pathogenicity [12, 14]. These tools typically utilise information from established variant datasets observed from biological samples of different extents of pathogenicity, to build classifiers using features which distinguishes pathogenic from nonpathogenic variants. While existing tools may demonstrate a good level of sensitivity and specificity in classifying pathogenicity, this does not necessarily mean these predictions offer accurate indications of protein-level impact of variants, partly a consequence of limited availability of variants which truly damage the stability and/or activity of the affected protein for predictors to be trained on. Since DMS experiments typically measure variant impact under a background of selection at the cellular level [49], learning on DMS data offers a route to provide annotation of variant impact which may not be directly associated with pathogenicity. Existing variant pathogenicity predictions are effective in assisting routine genetic diagnoses; however, given the advancement of sensitive detection of ultra-rare variants in the genome [44], DMS data can potentially offer an additional filter to identify truly damaging variants ignored by conventional pathogenicity predictors. We have encouraging results from our gradient boosting classifiers (Figures 5,6). Here, in comparison with the common somatic mutational signatures which describe the causes of mutations (and therefore their presence is directly indicative of the activity of the given mutational process, see e.g. [50]), the signature for damaging protein variants (Figure 1B) which we identify here can be coupled with contextual information about the environment around which mutations are found to occur (e.g. conservation, solvent accessibility etc.) to achieve an accurate elucidation of the likely variant impact. Our predictors show that even experimental data on a handful (≈ 10) of proteins can already yield a reasonably accurate variant impact predictor which learns generalised features that can be applied on unseen proteins, at an accuracy similar to Envision [36], which is also trained on DMS data to predict variant impacts. These DMS-based machine learning approaches can (a) impute variant effects for proteins which have not been subjected to experimental DMS, and (b) inform more economical mutant library designs if the goal of the DMS experiment is to screen for “fragile points” of the protein which are important to protein stability/activity; here our results suggest a proline scanning experiment is particularly revealing of variant impact, as evidenced by its heightened contribution to the predictor performance (Figure 6C); Sruthi and Prakash [37] have suggested that conversions to alanine, histidine and asparagine constitute a minimal set of mutants to assay experimentally, and impute effects of the remaining variants with reasonable accuracy. More systematic evaluations on a larger and more representative set of experimental DMS data will inform a rational design of mutant libraries to maximise utility of the data generated while saving costs.

The DNA mutational signature for damaging protein variants described in this work opens questions as to how much of our proteome is vulnerable to such damage. The conservation of the protein core, which is known to be under the slowest evolutuionary rates in comparison to other protein regions [43], necessitates the preservation of stability by maintaining polarity, hydrophobicity and packing arrangements of side chains in this region [51]. Retaining and/or accumulating amino acids which contribute to core stability could come with a potential cost to cellular fitness, if such evolutionary constraint implies accumulation of DNA motifs which could act as “substrates” for mutagenesis. A systematic investigation of the relationship between different mutagenic exposures and evolutionary conservation of functional sites warrant careful sequence and structural homology mapping. This could be a further important step in understanding the potential risks different mutagenic exposures could pose towards protein function and therefore their contribution in disease processes, across species of varying lifespan and living conditions.

## Acknowledgement

This research was supported by Croucher Foundation Hong Kong (https://croucher.org.hk/, scholarship to JCN), the Medical Research Council (https://mrc.ukri.org/, MR/L01257X/1 to FF) and the Biotechnology and Biological Sciences Research Council (https://bbsrc.ukri.org/, BB/T002212/1 to FF and JCN). The funders had no role in study design, data collection and analysis, decision to publish, or preparation of the manuscript.

## Methods

### Variant datasets from observational studies

In this analysis we analysed a total of 27 cancer types from The Cancer Genome Atlas (TCGA; [31]) database, taking as input the annotated TCGA Mutation Annotation Files (MAFs) available on Broad GDAC Firehose (https://gdac.broadinstitute.org/). Variants were filtered to group together variants mapped to consecutive genomic positions in order to distinguish variants occurring in doublet-base substitution (DBS) signatures. Variants were mapped to protein structures using ZoomVar [12], using UniProt identifiers as keys. See below (section “Mapping of coding sequences and protein structural data”) for a detailed description of parameters used to define structural regions. Amino acids were classified by their physicochemical properties into one of the following 5 groups: (i) aromatic (Y, W, F); (ii) negative (D, E); (iii) positive (R, K, H); (iv) hydrophilic (S, T, C, P, Q, N), and; (v) hydrophobic (G, A, V, L, M, I). These amino acid groups were considered in the analysis of the physicochemical impact of the damaging mutational signature (Figure 2A).

### Mapping of coding sequences and protein structural data

#### Protein structural localization

For each protein we obtained residue-level mapping of its core, surface and protein-protein interaction interface from the ZoomVar database [12]. In ZoomVar, protein positions are classified into protein core, surface and interacting interfaces. Briefly, core residues are defined to have a fractional solvent accessible surface area (SASA) less than 0.15 (calculated using POPS [35]); otherwise they are labelled as surface. Interfaces were annotated using POPSCOMP [52] by considering the change in solvent accessible surface area upon complex formation; residues were annotated to be in the interacting interface if such change was larger than 0.15 [53].

#### Enumerating coding sequences containing DNA mutational contexts

We obtained from UniProt its curated set of coding sequences (CDS) for the human proteome (UniProt Proteome UP000005640), and mapped, for each protein, the corresponding CDS sequence on a per-residue level. We then enumerated the percentage of residues within each CDS which could be altered by mutation within every possible trinucleotide motif. Since every position of the codon could be substituted to cause an amino acid change, trinucleotide mutational contexts could involve the codon immediately before (if codon position 1 was substituted) or after (if codon position 3 was substituted) the given amino acid position. We therefore considered, for each amino acid position, a 5-mer consisting of the triplet code for the position, plus one base 5’ and one base 3’ to the triplet code. Using these 5-mers we classified and counted whether each residue contains the named sequence motif.

### Quantification of variant/motif enrichment

For variant enrichment, we quantified such enrichment in a given protein structural region by calculating the following density metric *ω*:

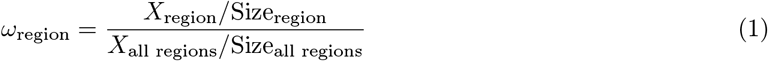

where

*X* is the number of mutations, and

“Size” refers to the number of amino acids belonging to the 3D structural region.

Similarly, the enrichment of motifs (Figure 2B) was calculated using the same formula as above, except that here *X*_region_ refers to the number of amino acids in the given 3D region that harbour the motif in question, and *X*_all regions_ refers to the analogous amino acid counts in all regions together.

### Deep mutational scanning (DMS)

#### Experimental DMS data

Experimental DMS datasets were downloaded from MaveDB [27]. We first considered three DMS datasets as case studies: (i) DMS on MSH2 (MaveDB accession 00000050-a-1) activity as readout [22]; (ii) DMS on PTEN with activity measured by a lipid phosphatase assay (MaveDB 00000054-a-1) [20]; (iii) DMS on PTEN with protein abundance as readout (MaveDB 00000013-a-1) [19]. These datasets were chosen as case studies because they were the two examples human proteins which (a) have available DMS data deposited on MaveDB (accessed 5 January 2021) and (b) contain known missense cancer driver variants.

#### Computational DMS data

We sought to obtain analogous DMS datasets computationally, by applying mutational impact predictors over full-length coding sequences of a comprehensive set of human proteins, with annotation of the mutational contexts of each assayed AA substitution. We considered all proteins with mutations observed in the TCGA dataset and took the best PDB match (see above) as the corresponding 3D structure as input to the mutational impact predictors. Due to potential one-to-many mappings between protein and transcripts, we prioritised transcripts with matched annotation between NCBI and EBI (from RefSeq MANE; https://www.ncbi.nlm.nih.gov/refseq/MANE/) in order to obtain unambiguous mapping of mutational contexts for each substitution. After this filtering, 6,918 unique Ensembl protein entries were subjected to *in silico* saturation mutagenesis using the rhapsody package [24]. Since the run also involved results from EVmutation [29] and PolyPhen-2 [28], these 3 predictors were considered alongside with rhapsody output. This dataset was considered in comparing the associations of different DNA trinucleotide motifs with damaging protein impact.

We used two additional DMS datasets outside of the analysis to identify the damaging mutational signature. First, results from the boostDM [26] algorithm were downloaded from https://www.intogen.org/boostdm/downloads. Second, predicted mutational effects on 6,938 human proteins were directly downloaded from EVmutation [29] (URL https://marks.hms.harvard.edu/evmutation/index.html). boostDM and EVmu-tation data were used in Figure 4 to define the variant landscape; EVmutation data were also used in training gradient boosting classifiers (see section “Gradient boosting classifiers to predict variant impact”).

#### Mapping DNA mutational contexts to amino acid substitutions

DMS data contain read-out/predictions for every amino acid position within a protein, mutating to every other 19 possible amino acids. A set of R functions, made available in the CDSMutSig package which accompanies this manuscript (see section “Code Availability”), was developed to map DNA mutational contexts to DMS measurements. We mapped Ensembl (version 86) proteins to their corresponding transcripts, obtained the coding sequence (CDS; genome build GRCh38), and extracted the DNA substitution as well as three-base sequence context surrounding the nucleotide substitution for each amino acid mutation. Amino acid substitutions which did not arise from a single-nucleotide variant (SNV) were removed.

#### Testing association between mutational impact and sequence context

All mutational impact scores were harmonised such that more negative/smaller values indicate more damaging variant impact. To examine the association between mutational impact and sequence contexts, the following linear model was considered:

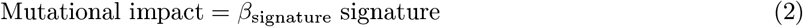

where

“Mutational impact” refers to the impact scores obtained from computational/experimental DMS profiles, and “signature” corresponds to each of the 96 possible three-base mutational contexts.

Such model was fit onto each of the 6 DMS datasets (3 [MSH2 activity, PTEN activity, PTEN abundance] experimental DMS profiles and 3 [rhapsody, PolyPhen-2 and EVmutation] computational DMS predictions), using the R glm function. The magnitude and *p*-value of *β*_signature_ from the linear model were taken for analysis.

### Gradient boosting classifiers to predict variant impact

#### Datasets

Two datasets were acquired to build the classifiers. First, we obtained all experimental DMS datasets from MaveDB [27] which assayed human protein coding sequences (accessed 24 August 2021). These datasets were manually annotated to (a) re-scale the scores such that a lower score correspond to a more damaging impact, and; (b) remove those datasets which did not display a bimodal score distribution (and hence was not amenable to a binary classification problem as framed in the main text; see Results). A total of 10 experiments covering 9 proteins remained: APP, SUMO1, PTEN, MSH2, TP53, TPMT, CBS, NUDT15, HRAS. This dataset is hereafter referred to as the “DMSexp” dataset. The list of MaveDB accession numbers for these datasets is included in Supplementary Table S2. Second, EVmutation [29] scores covering 6,938 human proteins were used as a larger dataset to address for the limited scope of the DMSexp dataset. Hereafter this dataset is referred to as “EVmutation”. EVmutation was chosen over other *in silico* scoring systems considered in this work because of the fact that it depended on statistical potentials modelled on the protein multiple sequence alignment but not on scores imputed from machine learning algorithms, and therefore avoided accumulating errors on top of predictions from other machine learning architectures. The EVmutation scores accounting for epistatic effects (i.e. co-dependency between residue positions in a multiple sequence alignment) were considered. Only variants which reside at amino acid positions in a Protein Data Bank structure were mapped (using ZoomVar [12]) and retained. BRCA1 (for both DMSexp and EVmutation) and PTEN (for EVmutation only) variants were removed prior to training as experimental data covering these two proteins were designated for evaluating the predictors (see below). The final datasets consist of 20,352 (DMSexp) and 5,805,302 (EVmutation) variants respectively. See Supplementary Table S1 for details.

#### Data processing

DMSexp and EVmutation scores were converted into binary labels (damaging/not damaging) in the following manner: for DMSexp, we performed K-means clustering to separate the data points into two clusters. Each DMS experiment was considered separately to avoid the effects of differing numeric scales of the scores between experiments. For EVmutation, we labelled those variants with score < −5 as “damaging” (and otherwise as “not damaging”); the original EVmutation paper [29] illustrated that this cut-off separates disease variants from common ExAC [54] variants. All variants were annotated for a set of mutational features:

1. amino acids (both wild-type and mutant) were encoded in the one-hot manner resulting in 20 (wild-type) + 20 (mutant) = 40 features.
2. DNA mutational signatures were considered as trinucleotide motif around the substitution, extracted using the functionalities implemented in CDSMutSig accompanying this manuscript (see section “Code availability”). The motif constituted, as one-hot encoding, a total of 14 features: 4 features for position -1 (A, T, C, G), 6 features for position 0, i.e. the position of nucleotide substitution (C*>*A, C*>*G, C*>*T, T*>*A, T*>*C, T*>*G), and 4 features for position +1 (A, T, C, G). Substitutions which remove a guanine or adenine were mapped to the opposite strand.
3. Sequence conservation was considered at the nucleotide level for positions -1, 0 and +1 with respect to the substitution, by mapping the phyloP [34] scores using the R GenomicScores package. phyloP scores for the three positions constistute 3 separate features.
4. Residue solvent accessibility annotated as Q(SASA) (quotient SASA, i.e., SASA relative to overall surface area) calculated using the POPS [35] package.

These annotations result in a 58-element (= 40 + 14 + 3 + 1) feature vector for each mutation. We divided DMSexp and EVmutation respectively into training and testing set as follows: for DMSexp, 20% of variants were randomly sampled and set aside as testing set; the remaining 80% was resampled using SMOTE [55] (implemented in the python imblearn package, version 0.8.1) to improve the balance between binary labels by over-sampling the depleted class. For EVmutation, we noted that the dataset was much larger and also fairly balanced (score = −5 was close to the median score = −5.3616) so no SMOTE sampling was performed; 1% of variants were randomly sampled and withheld as the test set, and the remaining 99% constitute the training set.

#### Predictor architecture, parameters and training

Gradient boosting classifier was chosen due to good support for methods to interpret feature importance (e.g. using Shapley additive explanations, or SHAP [39]), and previous success in other variant impact prediction tasks (e.g. boostDM [26]). We used the gradient boosting classifier implemented in the scikit-learn (version 0.24.2) python package. A grid search using 10-fold cross-validation on the DMSexp training data was performed to identify the following set of hyperparameters: learning rate = 0.1, maximum depth of individual classifiers = 6, number of boosting stages = 500, proportion of samples to fit individual classifiers = 0.7. All classifiers were built using scikit-learn with this set of hyperparameters, on the different feature combinations (Figure 5) on the training sets from DMSexp and EVmutation.

#### Performance evaluation

Classifiers were evaluated first on the holdout test sets respectively for DMSexp and EVmutation, and on the following datasets covering proteins unseen from training: (1) an experimental DMS dataset on the BRCA1 protein [21] was tested on both DMSexp and EVmutation classifiers; (2) the PTEN abundance and activity DMS datasets described in the section “Deep mutational scanning” (see above) was tested on the EVmutation classifiers. All variants in these independent test sets were annotated to obtain the input feature matrices in the same manner described above. The following metrics were calculated during evaluation: receiver operating characteristic (ROC) curve, ROC area-under-curve (ROC-AUC), accuracy, and F1-score. Additionally, to generate head-to-head comparisons with Envision [36] (which was trained also on experimental DMS data but framed as a regression task), experimental DMS data on *n* = 5 proteins (TPK1, LDLRAP1, UBE2I, SRC, HMGCR) were collected from MaveDB [27], the variant scores normalised using the workflow implemented in the deepscanscape R package [56]. All five proteins were unseen by the DMSexp predictor. DMSexp was used to predict these variants, and the predicted damaging probability (i.e. probability that the label is 1 [damaging]) is compared against the experimental DMS scores using Pearson’s correlation. Envision predictions on these proteins were downlaoded from Envision (URL: https://envision.gs.washington.edu/shiny/envision_new/) on 31 January 2022, and the analogous correlation metric was calculated between the Envision-predicted scores and the experimental DMS scores. Feature importance of the DMSexp and EVmutation predictors was calculated as the absolute Shapley additive explanation (SHAP) values (expressed as |SHAP| in Figure 6; the higher this value, the higher importance it contributes towards the predictions) using the python shap package (version 0.40.0) evaluating on the holdout test data for all classifiers. Absolute SHAP values for individual one-hot-encoded features were summed together, separately for each prediction made, to quantify importance of the overall feature category (Figure 6B).

### Statistics and data visualisation

All statistics were performed under either the R statistical computing environment (version 4.0.2). Linear models were fitted using the glm function in R. The variant impact predictors were generated using python packages as described in the section “Gradient boosting classifiers to predict variant impact”. All plots were generated using plotting functionalities in base R and the ggplot2 package [57].

## Code availability

The CDSMutSig R package provides functionalities to map between protein and DNA coding sequences and obtain trinucleotide mutational contexts for any amino acid mutations. This package is available at https://github.com/josef0731/CDSMutSig. All analysis code can be found at https://github.com/josef0731/mutational-dark-matter.

## Data availability

All data files generated for this analysis is available at this Zenodo repository: https://doi.org/10.5281/zenodo.4726168. A web application is also available for users to query somatic variants analysed in this paper, view structural annotations and map these variants to 3D structures. This server is available at fraternalilab.kcl.ac.uk/ZoomVar/ZoomvarSomatic/.

