## Supplementary Materials for "The “dark matter” of protein variants carries a distinct DNA signature and predicts damaging variant effects"

### Supplementary Note 1. Recurrent mutations attributable to mutational signature case studies.

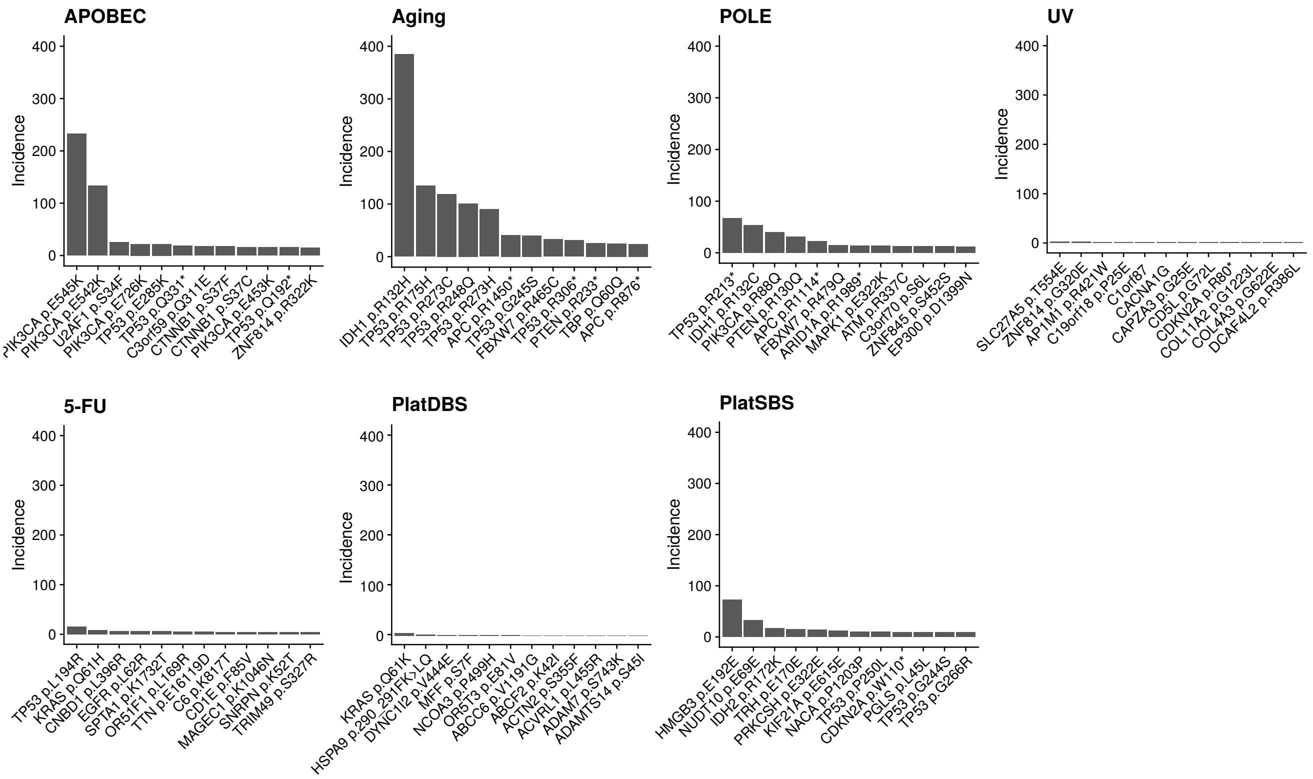


Figure N1.1: Top most frequent recurrent substitutions in the TCGA dataset occurring within the sequence context of each mutational signature in this paper. The "APOBEC" panel is identical to that shown in Figure N1.3D (see below) except that a different range in the vertical axis is used here.

We investigate whether there are individual positions harbouring recurrent variants observed in tumour samples which could be attributed to the mutagenic processes which we select as case studies in the main text (see Figure 2 in the Results section). As introduced in the main text, the attribution of individual mutations to specific mutagenic processes remains a challenge in mutational signature extraction tools. Indeed, in mutation data from The Cancer Genome Atlas (TCGA) database, we do not find recurrent hotspots occurring in DNA motifs attributable to the UV, 5-FU and Platinum signatures (Figure N1.1); there are well-characterised mutational hotspots occuring in the aging signature context, however these mutations do not appear to be associated with age of cancer onset (Figure N1.2), suggesting these could also be coincidental associations.

However, we find recurrent mutations for the POLE and APOBEC signatures which are conceivably associated with these aetiologies. Firstly, we observe the mutation PTEN p.R130Q which is present almost exclusively in POLE-mutated hypermutated tumours (Figure N1.3A). Arg130 is a buried position close to the active site of the PTEN phosphatase domain (Figure N1.3B) [4]; the R→Q mutation is predicted to be destabilising to the protein (Figure N1.3C), suggesting its possible loss-of-function effect, although its effect is not extreme relative to other amino acid substitutions arising from point mutations at this position. For the APOBEC signature, we notice that four mutations in PIK3CA are amongst the most frequent substitutions

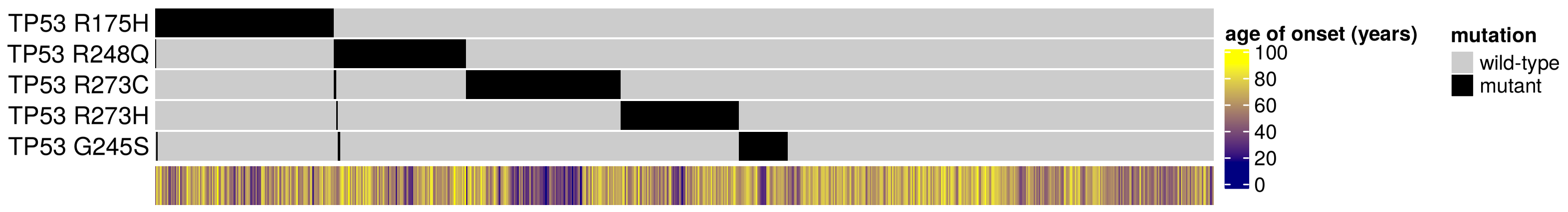


Figure N1.2: Occurrence of selected *TP53* mutations and the age of cancer onset in the TCGA dataset.

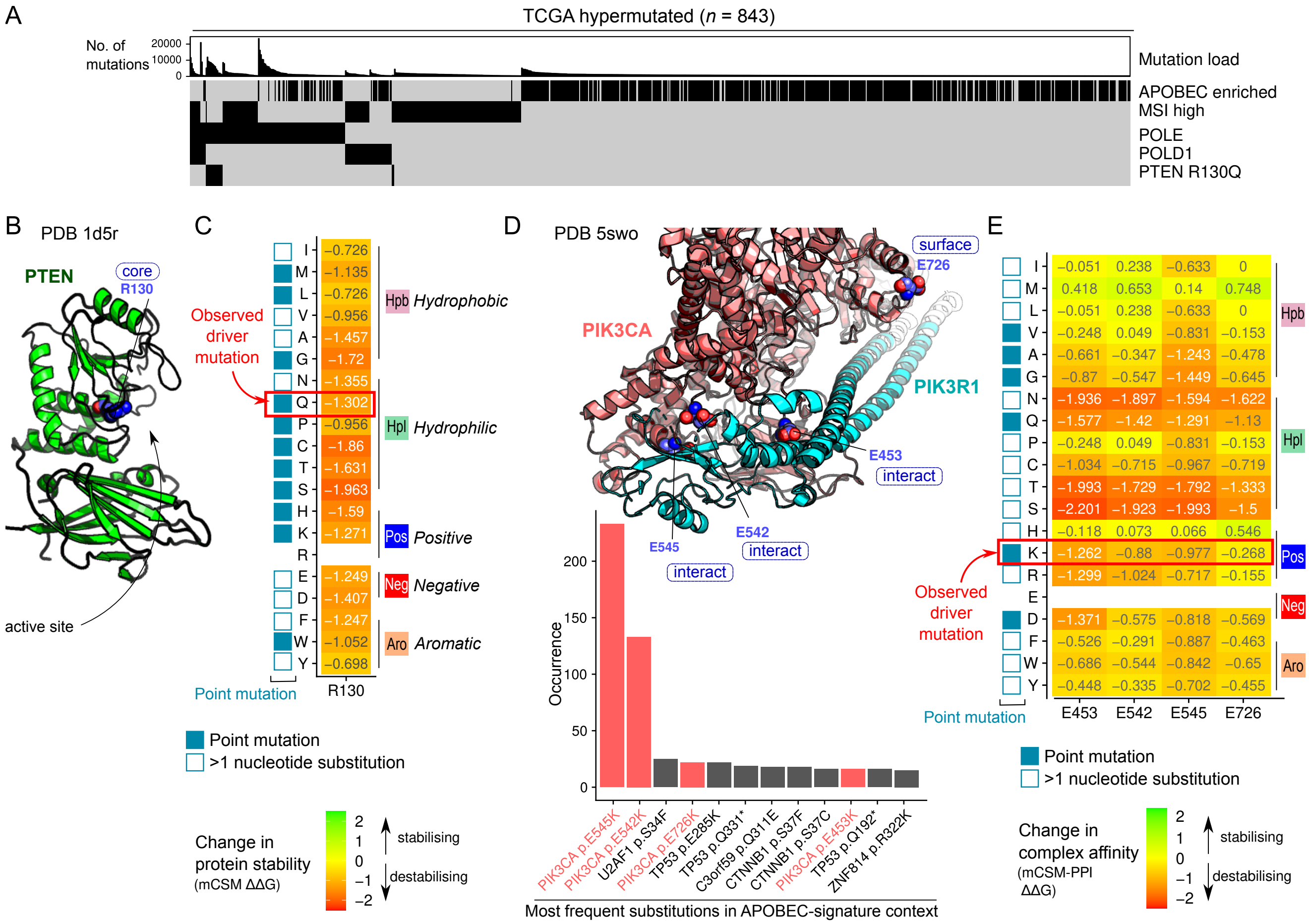


Figure N1.3: Mapping somatic mutagenic processes to protein structures. (A) Occurrence of molecular alterations across $n = 843$ TCGA hypermutated cases (arranged in columns). APOBEC signature enrichment is calculated as described [1]. MSI (Microsatellite instability) status is taken directly from TCGA clinical annotations. Sequence alterations in POLE, POLD1 and the PTEN p.R130Q mutations are indicated. (C) p.R130Q mutation mapped onto the PTEN structure (Protein Data bank [PDB] 1d5r). (D) Change in protein stability predicted using mCSM [2] for the PTEN Arg130 position. Points mutations and amino acid (AA) changes arising from more than 1 nucleotide substitution are indicated next to every assayed AA change. (E) (bottom) Most frequent substitutions in the APOBEC signature. PIK3CA mutations are highlighted in red. (top) Structural view of PIK3CA (red) in complex with PIK3R1 (cyan) (PDB 5swo). Positions with recurrent mutations are highlighted. (F) Change in complex affinity predicted using mCSM-PPI [3].

in the APOBEC signature context (Figure N1.3D); two of these (p.E545K and p.E542K) are well-documented mutational hotspots which are linked to elevated APOBEC mutagenesis [5]. These variants reside at the interface between PIK3CA and its regulatory partner PIK3R1 (Figure N1.3D) [6], and are predicted to be mildly destabilising to this protein complex (Figure N1.3E) [3]. In summary, these analyses map the protein structural impact of different mutagenic processes, and link these mutations to the biophysical basis of their possible alteration towards function. Interestingly, the observed recurrent mutations are often not the substitution which could theoretically confer the most dramatic impact; the most damaging amino acid changes often require the rarer mutational event of tandem nucleotide substitutions (Figure N1.3C,E). This suggests a degree of protection from the rule-book of codon usage and sequence preferences of mutagenic processes, in that highly damaging protein variants require successive point mutations to occur.

### Supplementary Note 2. Features of mutations in different protein structural regions.

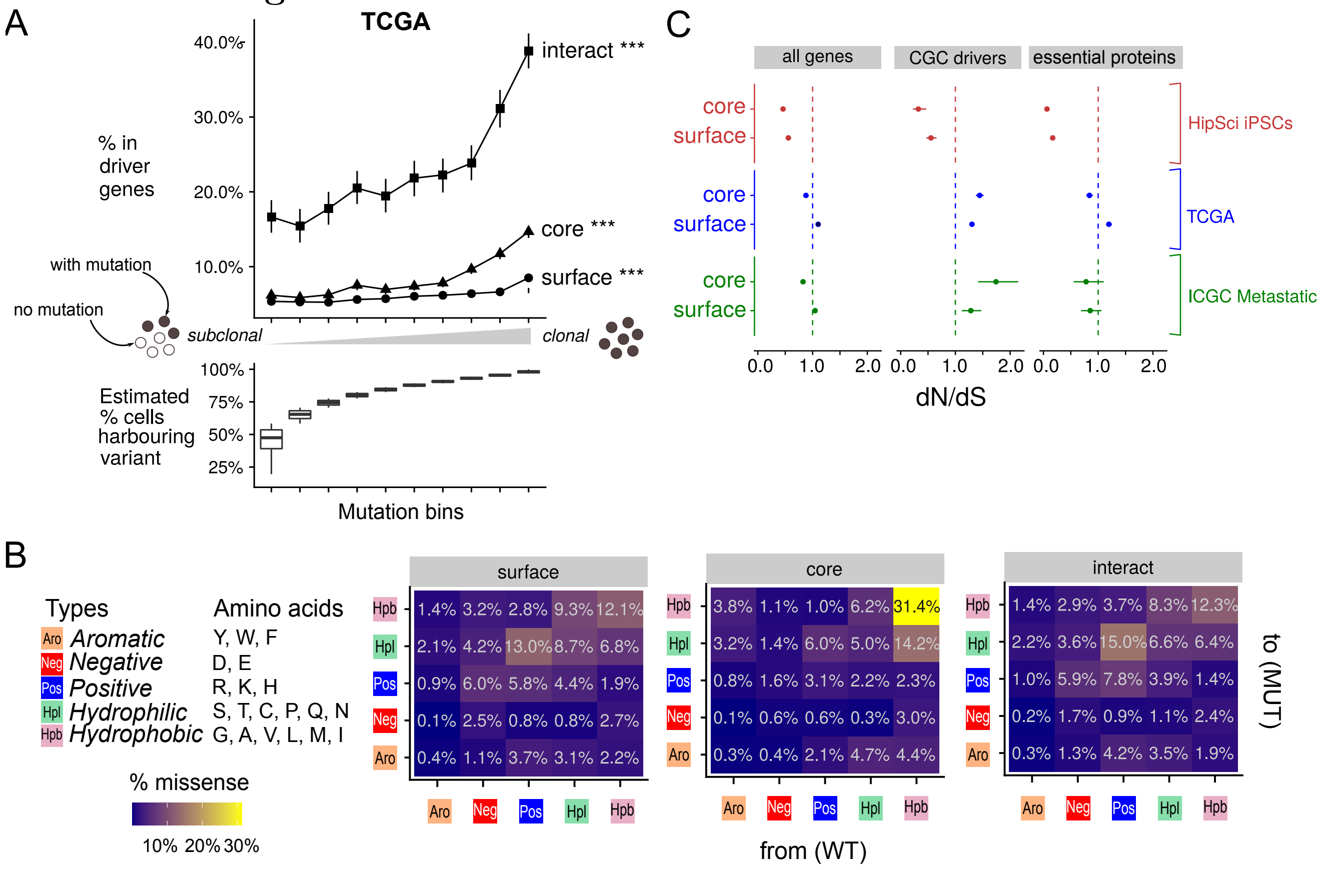


Figure N2.1: Protein structural localisation is indicative of driver status, clonality and selection pressure. (A) The percentage of mutations that localise to a COSMIC Cancer Gene Census (CGC) driver gene across the spectrum of clonality (measured by the estimated proportion of cells harbouring such mutations, see Supplementary Methods), considered separately protein surface, core and interface. Error bars depict 95% confidence interval estimated by bootstrapping. Trends are evaluated using the Jonckheere-Terpstra Test; ***, $p = 0.001$. (B) Distribution of variants in essential proteins (see Supplementary Methods) which change from one amino acid (AA) type to another in surface, core and interacting interface. The wild-type (WT) residue type is depicted on the horizontal axis and the mutant (MUT) residue on the vertical axis. The proportion of variants for each transition type is noted. (C) dN/dS comparison of protein surface and core, calculated using the dNdScv package [7]. Error bars show 95% confidence intervals. As controls the same calculation is performed on two gene sets: Cancer Gene Census (CGC) drivers and essential proteins. Table N2.1 below contains the values visualised here.

Here we characterise the features associated with variants observed in tumour samples falling into protein surfaces, cores and interacting interfaces, both at the molecular (the types of physicochemical changes impart on the affected protein) and cellular (frequencies within the tumour and selection pressure) levels. First, we observe that variants in protein core or interacting interfaces are more likely to reside in cancer drivers if they are more clonal (Figure N2.1A), i.e. occur earlier in tumour evolution and therefore are more frequent across the population of tumour cells. This indicates that not only protein structural localisation of variants indicates driver status (as previously reported, e.g. [8]), but also a reflection of the clonality of mutations. We further observe, using Gene Set Enrichment Analysis (GSEA) [9], that clonal mutations at interacting interfaces perturb important signalling pathways and cell adhesion, while those at the core perturb proteins such as transporters and molecules responsible for binding ions and nucleotides (Figure N2.2).

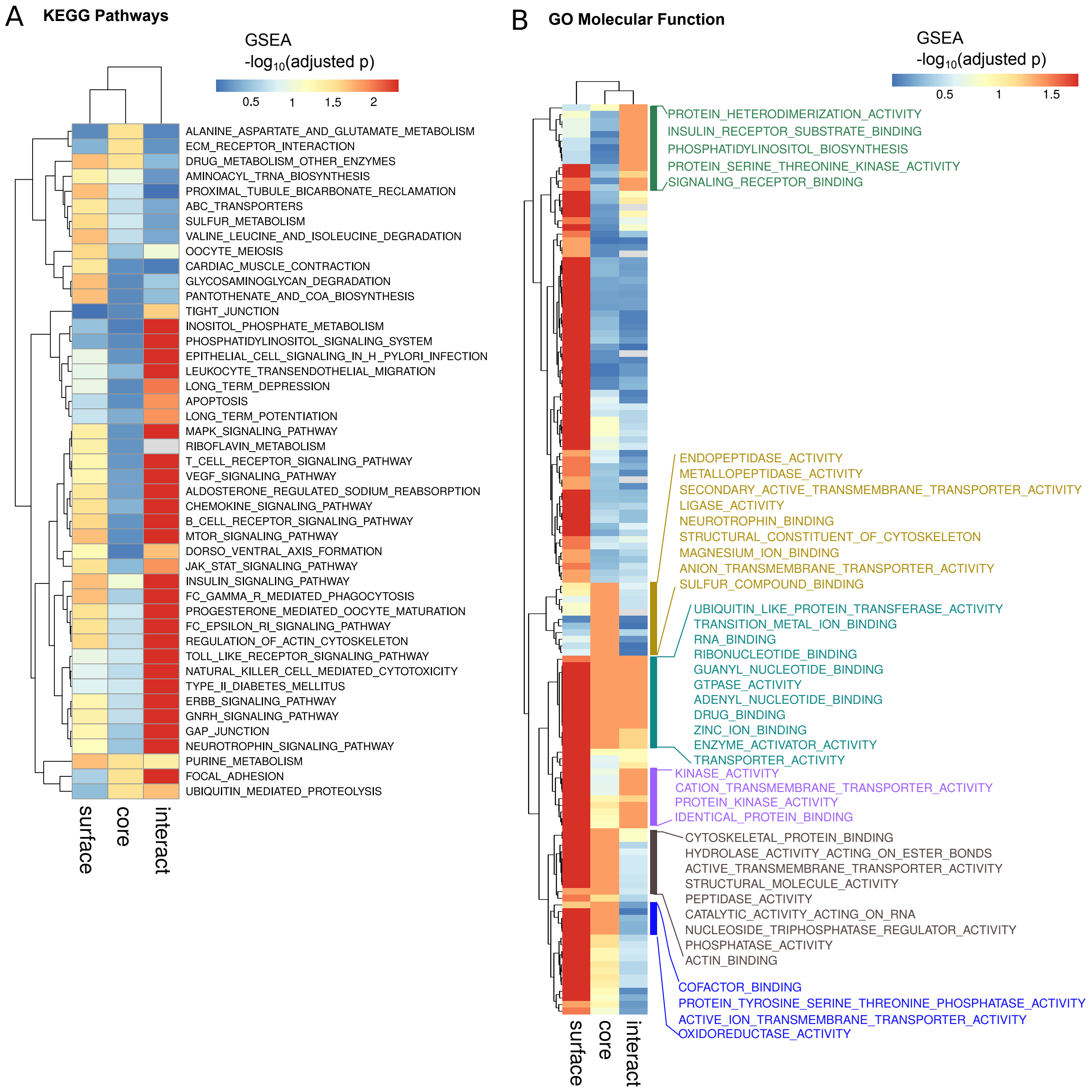


Figure N2.2: Gene Set Enrichment Analysis (GSEA) of clonal TCGA mutations in the protein surface, core and interacting interface. Mutations are ranked by their clonality (i.e. estimated proportion of tumour cells harbouring the given mutation) and the conventional GSEA procedure is applied to (A) KEGG Pathways and (B) GO Molecular Function. For (B), GO terms enriched for clonal mutations in core and interacting interfaces are highlighted.

Considering amino acid changes observed in tumours, substitution profiles in protein surface, core and interface reflect differences in solvent accessibility, with substitutions amongst hydrophobic amino acids much more common in the protein core (Figure N2.1B). On the other hand, changes from positive to polar amino acids are prominent in protein surfaces and interacting interfaces. The similarity between data from protein surfaces and interacting interfaces implies that the likelihood of transition between amino acid types are indications of solvent accessibility. The more restrictive nature of substitutions in the core is applicable both to a set of essential proteins whose removal would render cancer cells nonviable as assayed in publicly available gene knockdown screens [10, 11] (Figures S2 and N2.3), and generally to whole TCGA and COSMIC cohorts (Figure N2.4).

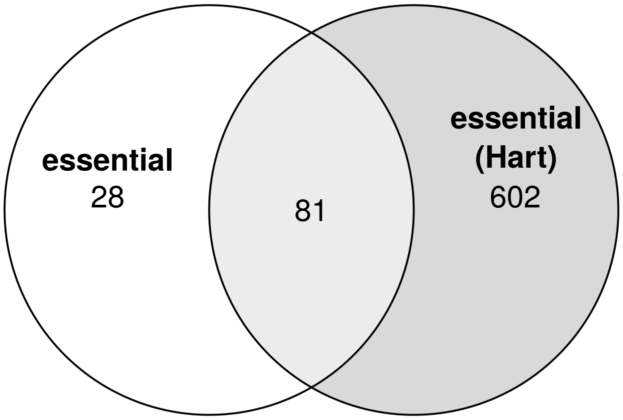


Figure N2.3: Overlap of the set of essential proteins defined as in Figure S2 and a published set obtained from [12].

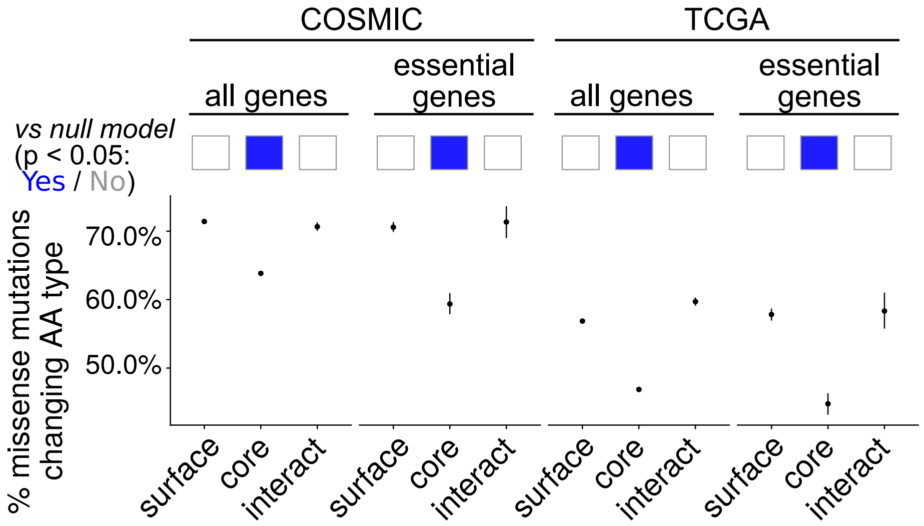


Figure N2.4: Proportion of variants changing amino acid types in COSMIC and TCGA datasets. The scheme to classify amino acid types is included in main text Figure 3C. We indicate also cases where significant difference is observed between the true statistic and null distributions where AA types are randomly assigned.

We investigate further the hypothesis that the core is subjected to greater purifying selection pressure in somatic evolution by explicitly calculating the dN/dS metric [7] which quantifies selection pressure, using datasets representing three different states in somatic evolution: (a) *in vitro*-cultured induced pluripotent stem cells (from the HipSci project [13]; see Figure S1 and Methods for details) (b) TCGA (mainly primary tumours) and (c) a dataset of metastatic cancers from the International Cancer Genome Consortium (ICGC) [14]. Comparing protein surface and core, we observe that protein core has a much more negative selection pressure compared to protein surfaces (Figure N2.1C) and Table N2.1; this trend is consistent across all three examined datasets. As controls, we perform the same dN/dS calculations specifically on essential proteins; expectedly, for HipSci iPSCs which are non-cancerous, we observe negative selection at both protein surfaces and cores (Figure N2.1C). Signal for purifying selection is weaker in the TCGA and ICGC metastatic samples (dN/dS closer to 1, indicating neutral selection pressure), suggesting that such proteins are perturbed with variants as cells become cancerous. Importantly, dN/dS at the protein surface is higher than the core in all cases for such essential proteins. As a positive control, cancer drivers (COSMIC Cancer Gene Census [15]) show a positive selection pressure in cancerous states. Interestingly, this is the only scenario where dN/dS at the core is higher than that at protein surface in the cancerous states, consistent with positively selected mutations in tumour suppressor genes which preferentially reside in the protein core [16]. These results show a robust difference in selection pressure and mutability of protein surfaces and cores, which resembles previous data observed in species evolution [17]; in the case of somatic variants the same difference also applies, linking protein structural localisation to selection. The only exception is in cases where core mutations contribute proliferative advantage which overrides purifying selection arising from damaging protein activity and stability. The clonality analysis supports the importance of 3D structural properties in modulating the intratumoural frequency of somatic mutations.

| Cohort | Genes | dN/dS Overall | | dN/dS Surface only | | dN/dS Core only | |
|---|---|---|---|---|---|---|---|
| HipSci iPSCs | all | 0.578 | [ 0.568 - 0.587 ] | 0.557 | [ 0.538 - 0.577 ] | 0.460 | [ 0.430 - 0.491 ] |
| HipSci iPSCs | CGC drivers | 0.550 | [ 0.506 - 0.597 ] | 0.554 | [ 0.468 - 0.655 ] | 0.324 | [ 0.224 - 0.469 ] |
| HipSci iPSCs | essential genes | 0.185 | [ 0.165 - 0.208 ] | 0.169 | [ 0.139 - 0.206 ] | 0.064 | [ 0.042 - 0.097 ] |
| ICGC Metastatic | all | 1.004 | [ 0.990 - 1.020 ] | 1.046 | [ 1.015 - 1.078 ] | 0.825 | [ 0.784 - 0.867 ] |
| ICGC Metastatic | CGC drivers | 1.329 | [ 1.237 - 1.428 ] | 1.281 | [ 1.116 - 1.472 ] | 1.742 | [ 1.414 - 2.147 ] |
| ICGC Metastatic | essential genes | 0.912 | [ 0.819 - 1.015 ] | 0.854 | [ 0.687 - 1.060 ] | 0.779 | [ 0.552 - 1.100 ] |
| TCGA | all | 1.049 | [ 1.045 - 1.053 ] | 1.102 | [ 1.094 - 1.110 ] | 0.878 | [ 0.868 - 0.889 ] |
| TCGA | CGC drivers | 1.256 | [ 1.235 - 1.278 ] | 1.302 | [ 1.262 - 1.343 ] | 1.444 | [ 1.375 - 1.516 ] |
| TCGA | essential genes | 1.095 | [ 1.069 - 1.121 ] | 1.196 | [ 1.142 - 1.252 ] | 0.840 | [ 0.782 - 0.903 ] |

Table N2.1: Selection pressure calculated on different sets of genes in HipSci iPSCs, TCGA and ICGC metastatic cases. Considered separately for overall dN/dS and specifically for surface and core only. Calculated using the dNdScv package [7]. Data visualised in Figure N2.1C.

### Supplementary Methods

This section covers methods for analyses presented in Supplementary Notes 1 and 2.

#### Variant datasets

**Cancer.** In additional to the TCGA data (see main text Methods), our analysis also considered somatic mutation data of all metastatic tumours from the International Cancer Genome Consortium (ICGC; [14]) database were downloaded from the ICGC data portal on 03/06/2019, by selecting only the tumour samples with the annotation of relapse type "`Distant recurrence/metastasis`". A total of 785 ICGC metastatic tumour samples were analysed.

**Induced pluripotent stem cells (iPSCs).** Whole-exome sequencing (WES) variant data were acquired from the HipSci project [13], a collection of induced pluripotent stem cell (iPSC) and skin fibroblast cell lines from donors, as a dataset representing cancer-free but proliferative condition. Only donors without disease annotations were considered. The Variant Call Format (VCF) files generated from mpileup variant calls for all such iPSC and skin fibroblast lines were downloaded, where access is publicly available from the European Nucleotide Archive (ENA, Project accession PRJEB7243). For cell lines where multiple vcfs were available, data generated from Illumina HiSeq 2000 was taken. To filter for somatic variants acquired during the course of cell line derivation and passage (see below), we only considered iPSC lines from donors for which WES data on fibroblasts (progenitors of the iPSC lines) were also available. Only variants marked "PASS" in the vcf FILTER field was included for downstream annotation and analysis. A total of 300 iPSC lines from 192 donors were analysed. Selection was quantified for this dataset to compare with the two (ICGC metastatic and TCGA) tumour datasets.

#### Variant processing

**Variant filtering and data mapping.** For consistency, only missense variants lying in exon regions were considered in the ICGC dataset, since TCGA and iPSC variants were called from WES data. These were extracted by intersecting (using bedtools [v2.25.0]) each vcf file with entries marked "exons" in the Ensembl (release 93, genome build GRCh37) Gene Annotation File. For the iPSC data, variant data from each iPSC line was compared with that profiled on its fibroblast progenitor, and filter for only variants unique to the iPSC lines but not found in the fibroblast (Supplementary Figure S1). For both cell types, variants localising to HLA genes were filtered away owing to their intrinsic highly polymorphic nature.

We calculated the mutational density per exome by normalising the total count of mutations for a given exome by its size (38 Mb was taken as an estimate [18]). Tumours with density $>$ 10 mutations per Mb were taken as hypermutated [19]. Finally, a density-based enrichment of mutational signatures was calculated following the method in [1] for calculating APOBEC-signature enrichment.

**Variant annotation.** To unify with the TCGA annotated MAFs, variant data from both the ICGC metastatic cohort and the HipSci iPSCs were annotated with gene and protein information using Oncotator (v1.9.9.0; [20]). Oncotator prioritises a set of transcripts with reading frames identical to UniProt sequences; these are directly taken forward for mapping these mutations to structure using ZoomVar. Additionally, for TCGA variants the frequency of variants (in terms of the percentage of tumour cells harbouring variants, also

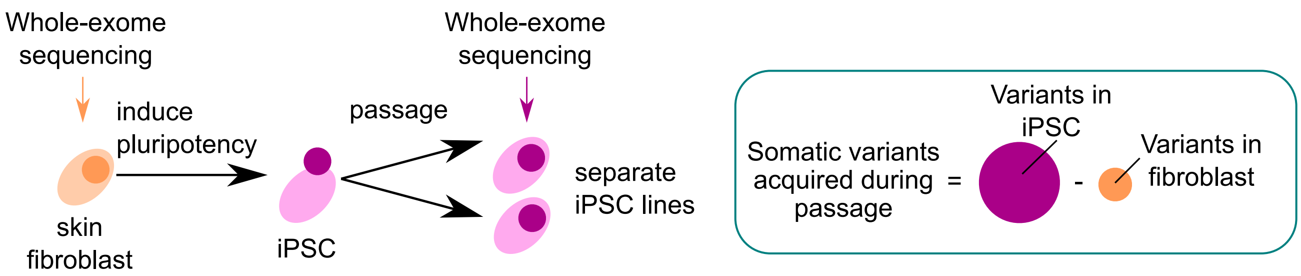


Figure S1: Schematic showing the filtering for variants acquired during iPSC derivation and *in vitro* culture of iPSCs.

known as “cancer cell fraction”) was estimated following the method described by McGranahan and colleagues [21].

**Annotation of protein stability change.** Using the ZoomVar database we annotated, for each TCGA variant with 3D structural coverage, the PDB entry with the highest sequence identity and lowest E-value match with the affected protein. We considered only those structures which harbours the same amino acid at the position in question as the wild-type. These variants were subject to variant impact prediction using the following tools: (i) the mCSM server [2] to predict the change in protein stability upon mutation; (ii) the rhapsody package [22] which predicts the probability that a given mutation is pathogenic, using results from PolyPhen-2 [23] and EVmutation [24], as well as coarse-grained network dynamics models defined on the input 3D protein structure [25].

#### Gene sets

**Cancer drivers.** The Cancer Gene Census (“CGC”, v86) was downloaded from the COSMIC database [15]. A gene set of cancer drivers was derived from the CGC by selecting only the “Tier 1” (documented relevance to cancer) genes, totalling to 574 genes.

**Essential proteins.** We utilised two data sources to obtain a list of essential proteins. First, a list of 109 essential proteins (see main text) was defined based on gene knockdown assays performed on cancer cell lines across cancer types. Both the CRISPR screen (AVANA project; [10]), and the RNAi screen (“Combined RNAi dataset” from the depmap project, based on short hairpin RNA [shRNA] libraries; see [26]), were downloaded from depmap (`https://depmap.org/`) and analysed. Both assays screened the impact of knocking down individual genes (CRISPR: 17,634 genes; RNAi: 17,309 genes) on the viability of cancer cell lines (CRISPR: 558 cell lines; RNAi: 712 cell lines) covering a total of 21 cancer types. This impact is represented by a “dependency score” akin to a $z$-score, where the more negative this score, the more dependent the cell line is to this particular gene (since its knockdown resulted in a significant decrease in viability, see Figure S2). We selected for genes demonstrating essentiality in all 21 cancer types, first calculating the mean dependency score for each gene in each cancer type, and then filtering for genes whose dependency score is in the top (most negative) 5% across all genes in all of the cancer types examined. Second, we collected another list of essential proteins defined using a different CRISPR knockdown dataset from Hart and colleagues [12]. This gene set was taken directly from `https://github.com/macarthur-lab/gene_lists`. We overlapped it with

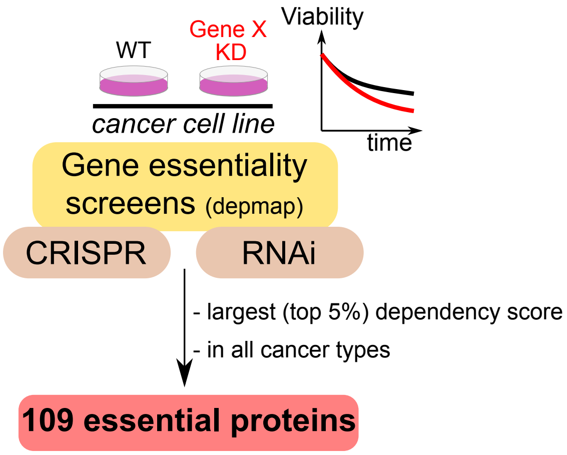


Figure S2: Defining proteins essential for cancer cell viability. A list of 109 essential proteins, consistent across cancer types, is defined using gene essentiality screen data, where cell viability in gene knockdowns (KD) is compared against the wild-type.

our own curated set in Supplementary Note 2.

**General pathways and molecular functions.** KEGG pathways and GO molecular function gene sets were acquired from MSigDB (v7.1) [9].

#### dN/dS calculations

dN/dS calculations were performed using the dNdScv package [7] was used here to quantify the dN/dS value for the iPSCs, TCGA and ICGC Metastatic datasets. Calculations were considered for protein surface and core by generating subsets of variants locating in the two regions and providing these two lists separately as input to dNdScv. Additional calculations on specific gene sets (cancer drivers, essential proteins) were performed by further filtering the dNdScv input by relevant gene symbols.

#### Supplementary Tables

| Dataset | No. of variants | No. of proteins |
|---|---|---|
| EVmutation | 5,805,302 | 6,671 |
| DMSexp | 20,352 | 9 |
| ThermoMutDB [27] | 11,199 | 448 |
| ThermoMutDB (Human only) | 2,298 | 134 |
| ProThermDB [28] | 16,113 | 414 |
| ProThermDB (Human only) | 3,586 | 116 |

Table S1: Number of variants and proteins in different datasets utilised in building gradient boosting classifiers (see main text Figures 5 and 6) in comparison to existing databases of thermostability impact of protein variants. Statistics for ThermoMutDB and ProThermDB were accessed at 13 December 2021.

| MAVEdb ID | Protein | Used in | Remarks |
|---|---|---|---|
| 00000001-a-4 | UBE2I | comparison with Envision | |
| 00000001-d-1 | TPK1 | comparison with Envision | |
| 00000035-a-1 | HMGCR | comparison with Envision | |
| 00000036-a-1 | LDLRAP1 | comparison with Envision | |
| 00000041-a-1 | SRC kinase domain | comparison with Envision | |
| 00000041-b-1 | SRC SH4 domain | comparison with Envision | |
| 00000001-b-1 | SUMO1 | training DMSexp | |
| 00000005-a-3 | CBS | training DMSexp | |
| 00000005-a-4 | CBS | training DMSexp | |
| 00000013-a-1 | PTEN | training DMSexp, fitting linear models | Protein abundance as readout |
| 00000013-b-1 | TPMT | training DMSexp | |
| 00000050-a-1 | MSH2 | training DMSexp, fitting linear models | |
| 00000054-a-1 | PTEN | fitting linear models | Protein activity as readout |
| 00000055-a-1 | NUDT15 | training DMSexp | |
| 00000057-a-1 | HRAS | training DMSexp | |
| 00000058-a-1 | APP | training DMSexp | |
| 00000059-a-1 | TP53 | training DMSexp | |

Table S2: List of MAVEdb experimental DMS datasets downloaded in this analysis.

| Test set | Training include core variants? | AA | MutSig | Conservation | Q(SASA) | DMSexp Accuracy | DMSexp F1 score | DMSexp ROC-AUC | EVmutation Accuracy | EVmutation F1 score | EVmutation ROC-AUC |
|---|---|---|---|---|---|---|---|---|---|---|---|
| holdout variants | ✓ | ✓ | ✓ | ✓ | ✓ | 0.825 | 0.714 | 0.868 | 0.734 | 0.696 | 0.808 |
| | ✓ | ✓ | ✓ | ✓ | | 0.792 | 0.649 | 0.827 | 0.709 | 0.665 | 0.774 |
| | ✓ | ✓ | ✓ | | ✓ | 0.783 | 0.690 | 0.837 | 0.720 | 0.679 | 0.789 |
| | ✓ | ✓ | ✓ | | | 0.601 | 0.505 | 0.645 | 0.686 | 0.633 | 0.741 |
| | ✓ | ✓ | | | | 0.596 | 0.500 | 0.637 | 0.686 | 0.633 | 0.741 |
| | ✓ | | ✓ | | | 0.578 | 0.442 | 0.588 | 0.593 | 0.473 | 0.614 |
| | | ✓ | ✓ | ✓ | ✓ | 0.712 | 0.567 | 0.749 | 0.697 | 0.601 | 0.778 |
| | | ✓ | ✓ | ✓ | | 0.724 | 0.460 | 0.681 | 0.670 | 0.521 | 0.763 |
| | | ✓ | ✓ | | ✓ | 0.653 | 0.587 | 0.717 | 0.671 | 0.553 | 0.750 |
| | | ✓ | ✓ | | | 0.619 | 0.354 | 0.576 | 0.631 | 0.420 | 0.729 |
| | | ✓ | | | | 0.628 | 0.366 | 0.583 | 0.631 | 0.420 | 0.729 |
| | | | ✓ | | | 0.644 | 0.282 | 0.564 | 0.552 | 0.064 | 0.604 |
| BRCA1 DMS | ✓ | ✓ | ✓ | ✓ | ✓ | 0.719 | 0.382 | 0.669 | 0.805 | 0.642 | 0.850 |
| | ✓ | ✓ | ✓ | ✓ | | 0.692 | 0.209 | 0.607 | 0.763 | 0.525 | 0.776 |
| | ✓ | ✓ | ✓ | | ✓ | 0.626 | 0.458 | 0.652 | 0.759 | 0.622 | 0.848 |
| | ✓ | ✓ | ✓ | | | 0.697 | 0.528 | 0.727 | 0.741 | 0.564 | 0.761 |
| | ✓ | ✓ | | | | 0.731 | 0.602 | 0.768 | 0.742 | 0.570 | 0.768 |
| | ✓ | | ✓ | | | 0.647 | 0.429 | 0.624 | 0.640 | 0.299 | 0.555 |
| | | ✓ | ✓ | ✓ | ✓ | 0.694 | 0.310 | 0.690 | 0.745 | 0.389 | 0.772 |
| | | ✓ | ✓ | ✓ | | 0.672 | 0.121 | 0.518 | 0.731 | 0.227 | 0.708 |
| | | ✓ | ✓ | | ✓ | 0.599 | 0.532 | 0.729 | 0.773 | 0.562 | 0.792 |
| | | ✓ | ✓ | | | 0.697 | 0.402 | 0.575 | 0.729 | 0.337 | 0.707 |
| | | ✓ | | | | 0.649 | 0.345 | 0.588 | 0.721 | 0.320 | 0.717 |
| | | | ✓ | | | 0.716 | 0.260 | 0.577 | 0.720 | 0.046 | 0.519 |
| PTEN stability DMS | ✓ | ✓ | ✓ | ✓ | ✓ | | | | 0.683 | 0.733 | 0.751 |
| | ✓ | ✓ | ✓ | ✓ | | | | | 0.615 | 0.672 | 0.720 |
| | ✓ | ✓ | ✓ | | ✓ | | | | 0.665 | 0.678 | 0.764 |
| | ✓ | ✓ | ✓ | | | | | | 0.683 | 0.670 | 0.744 |
| | ✓ | ✓ | | | | | | | 0.665 | 0.657 | 0.742 |
| | ✓ | | ✓ | | | | | | 0.502 | 0.375 | 0.541 |
| | | ✓ | ✓ | ✓ | ✓ | | | | 0.638 | 0.664 | 0.723 |
| | | ✓ | ✓ | ✓ | | | | | 0.611 | 0.598 | 0.706 |
| | | ✓ | ✓ | | ✓ | | | | 0.606 | 0.549 | 0.726 |
| | | ✓ | ✓ | | | | | | 0.566 | 0.360 | 0.704 |
| | | ✓ | | | | | | | 0.575 | 0.390 | 0.701 |
| | | | ✓ | | | | | | 0.489 | 0.066 | 0.510 |
| PTEN activity DMS | ✓ | ✓ | ✓ | ✓ | ✓ | | | | 0.616 | 0.586 | 0.820 |
| | ✓ | ✓ | ✓ | ✓ | | | | | 0.616 | 0.597 | 0.798 |
| | ✓ | ✓ | ✓ | | ✓ | | | | 0.716 | 0.641 | 0.815 |
| | ✓ | ✓ | ✓ | | | | | | 0.726 | 0.617 | 0.787 |
| | ✓ | ✓ | | | | | | | 0.712 | 0.605 | 0.788 |
| | ✓ | | ✓ | | | | | | 0.626 | 0.353 | 0.590 |
| | | ✓ | ✓ | ✓ | ✓ | | | | 0.672 | 0.618 | 0.771 |
| | | ✓ | ✓ | ✓ | | | | | 0.721 | 0.623 | 0.782 |
| | | ✓ | ✓ | | ✓ | | | | 0.719 | 0.560 | 0.750 |
| | | ✓ | ✓ | | | | | | 0.712 | 0.386 | 0.746 |
| | | ✓ | | | | | | | 0.709 | 0.402 | 0.745 |
| | | | ✓ | | | | | | 0.681 | 0.068 | 0.556 |

Table S3: Performance of gradient boosting classifiers trained on different features, tested on distinct test datasets. The Accuracy, F1 score and ROC-AUC metrics are listed. See Results and Figures 5 and 6 of the main text for details.

#### Supplementary Figures

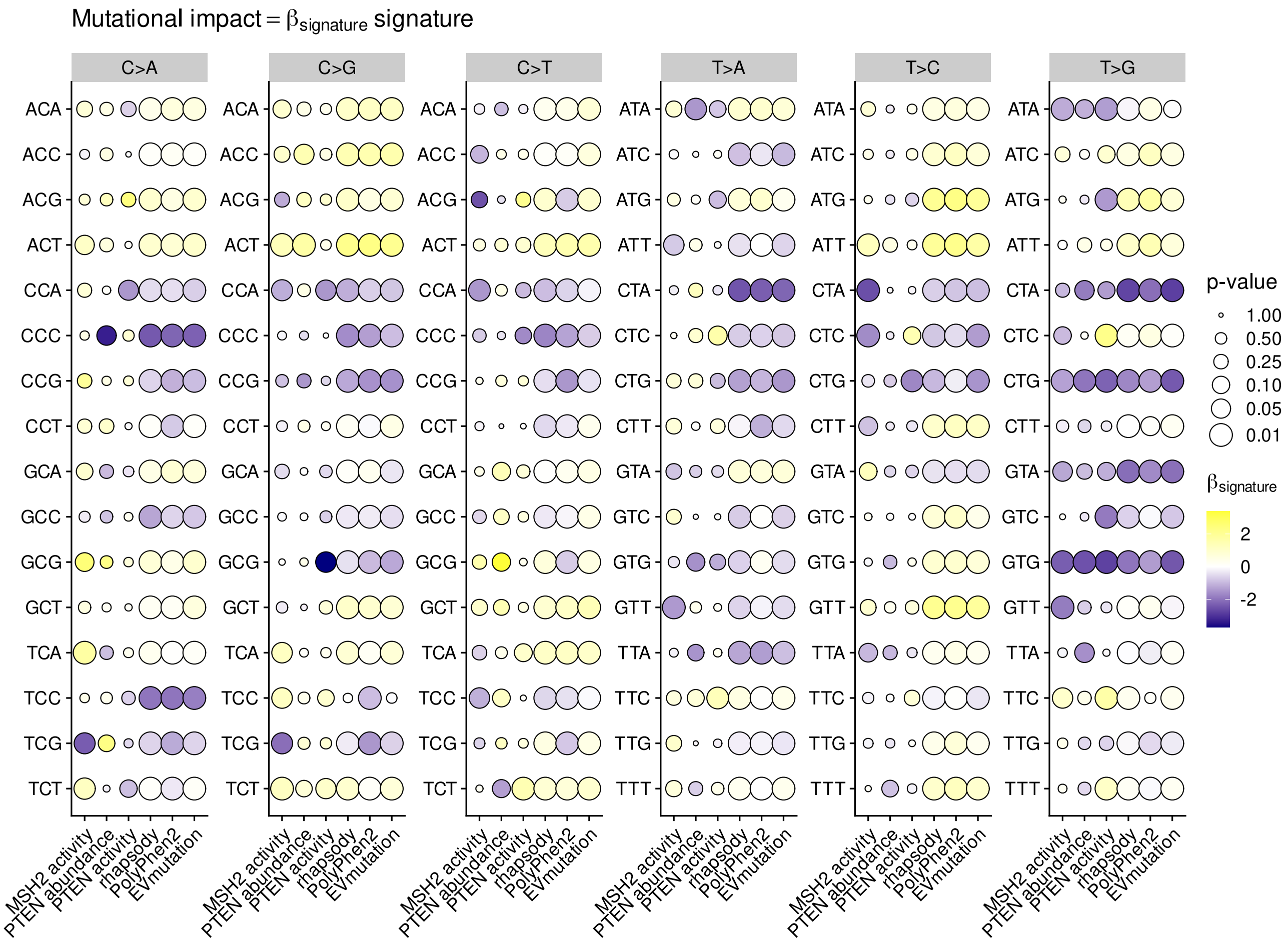


Figure S3: Association of DNA motifs to damaging impact. Identical to main text Figure 3C, here the magnitude and statistical significance of the coefficients of each context ($\beta_{\text{signature}}$) are visualised. All 96 DNA trinucleotide motifs used to define trinucleotide mutational signatures are included here. See Table S4 (XLSX file) for underlying data.

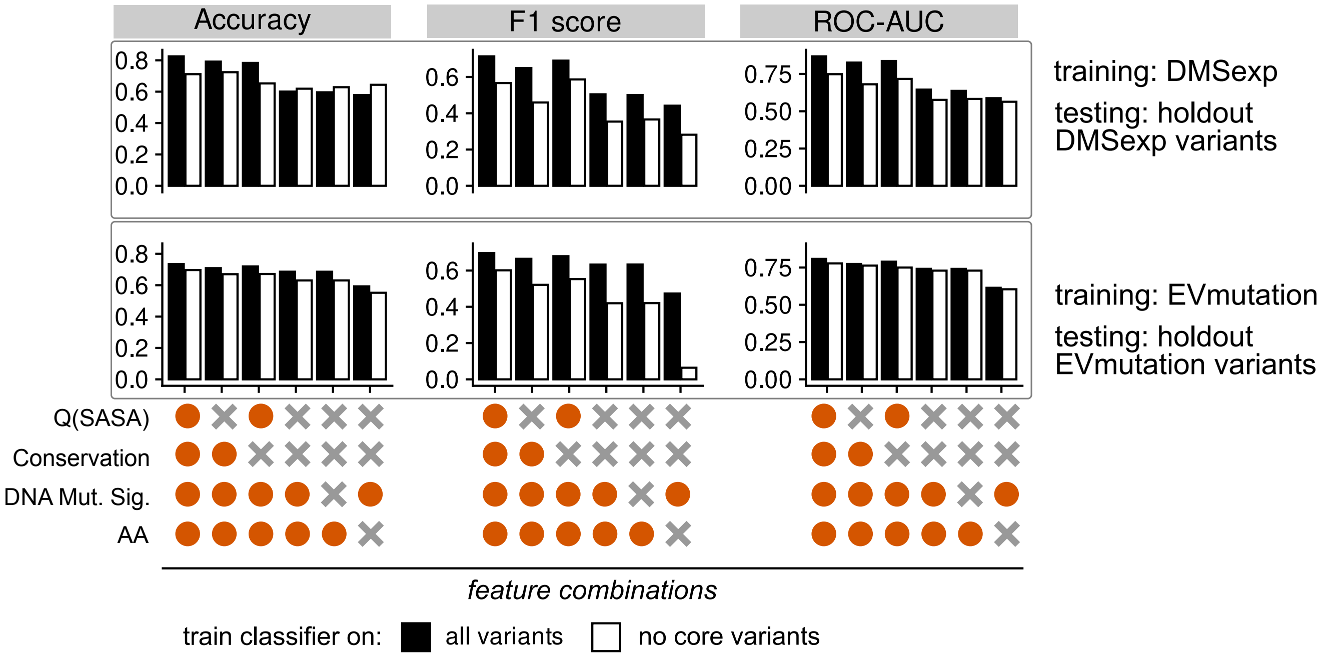


Figure S4: Performance of gradient boosting classifiers trained on DMSexp (top) and EVmutation (bottom) data on respective holdout variants as test set. Performance is compared between the "full" model including all variants (black bars) against a version where protein core variants (those with Q(SASA) $< 0.15$ are removed as a proxy of removing variants in the mutational "dark matter" (white bars).

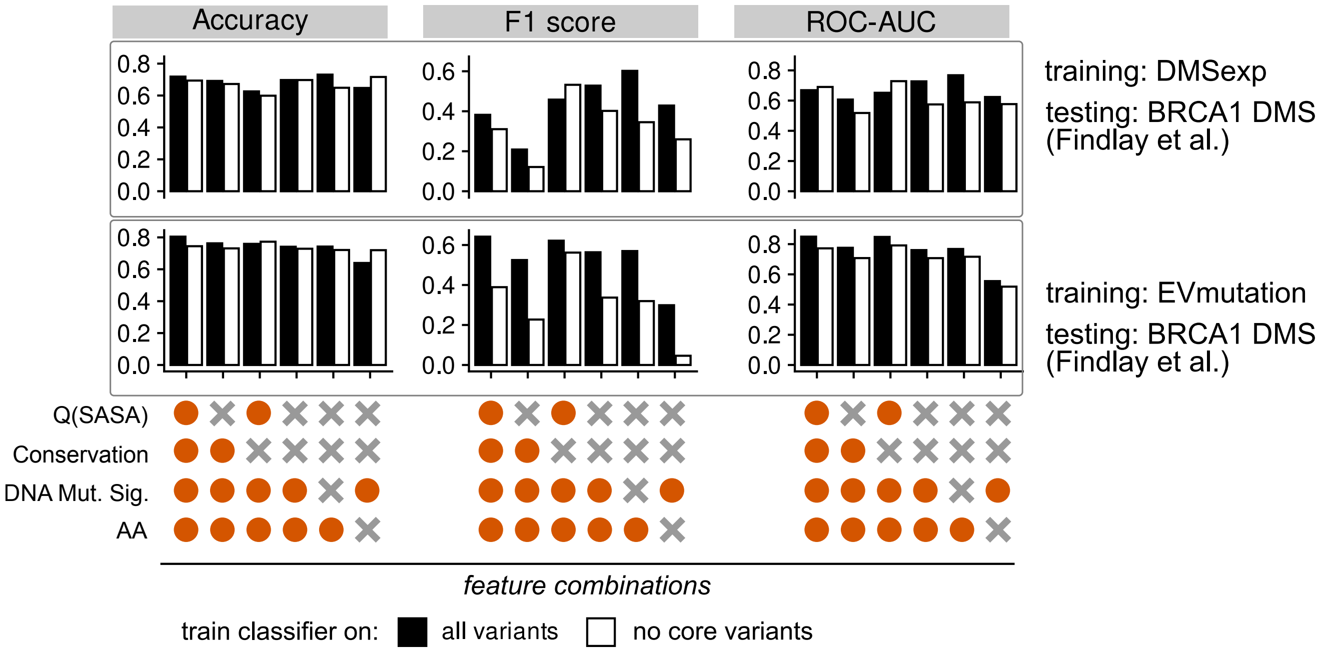


Figure S5: Performance of gradient boosting classifiers trained on DMSexp (top) and EVmutation (bottom) data on a BRCA1 DMS experimental dataset [29]. Performance is compared between the "full" model including all variants (black bars) against a version where protein core variants (those with Q(SASA) $< 0.15$ are removed as a proxy of removing variants in the mutational "dark matter" (white bars).

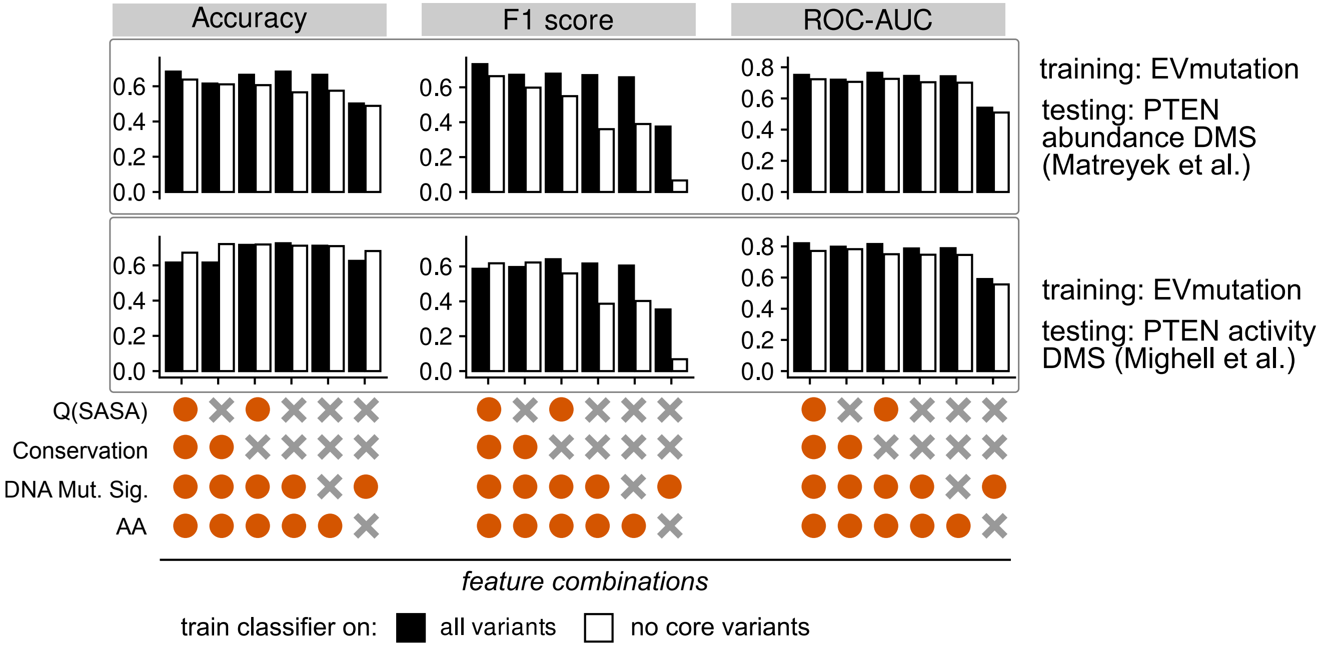


Figure S6: Performance of gradient boosting classifiers trained on EVmutation data using, as test set, DMS data collected on PTEN using protein abundance [30] (top) or activity [31] (bottom) as readout respectively. PTEN variants have been removed from the EVmutation data prior to training. Performance is compared between the "full" model including all variants or a version where protein core variants (those with Q(SASA) $< 0.15$ are removed as a proxy of removing variants in the mutational "dark matter" (white bars).
